# A benchmark of DNA methylation deconvolution methods for tumoral fraction estimation using DecoNFlow

**DOI:** 10.1101/2025.11.27.688590

**Authors:** Edoardo Giuili, Sofie Van de Velde, Sam Kint, Maísa R Ferro dos Santos, Lotte Cornelli, Sofie Roelandt, Kathleen Schoofs, Renske Imschoot, Ruben Van Paemel, Leander Meuris, Celine Everaert, Katleen De Preter

**Author notes:** Corresponding author: Katleen De Preter.

## Abstract

In cancer patients, circulating cell-free DNA (cfDNA) is released into body fluids from both healthy and cancer cells. The proportion of tumor-derived cfDNA serves as a surrogate marker of tumor burden allowing disease monitoring. Tumoral cfDNA can be distinguished based on patient specific tumoral mutations or using more general tumor specific DNA methylation patterns, that are preserved on tumoral cfDNA. DNAm profiling of cfDNA thus enables non-invasive cancer detection and monitoring. However, accurately determining tumour fractions remains challenging due to the heterogeneous mixture of cfDNA sources in body fluids. Computational DNAm deconvolution methods address this by inferring cell-type contributions either with or without reference methylomes. While several tools exist and multiple benchmarking studies have been performed, none have specifically evaluated the sensitivity and accuracy of tumour-fraction estimation in cfDNA-focused contexts. Here, we benchmarked 10 reference-based and 2 reference-free DNAm deconvolution tools using 3,690 in silico mixtures spanning multiple tumour types, different bisulfite-based sequencing strategies and several sequencing depths. Overall, CelFiE showed the most accurate tumour-fraction estimation across the different conditions. Interestingly, reference-free methods demonstrated superior sensitivity for tumour detection, but consistent over-estimation of tumoral fraction. We further observed that sequencing depth strongly affects performance until sufficient saturation is achieved. To enable reproducible evaluation and tool selection within this benchmark, we developed DecoNFlow, a scalable Nextflow pipeline integrating 12 deconvolution tools and 3 marker selection methods, making it the most comprehensive pipeline for sequencing-based deconvolution up to date. Together, our findings provide practical guidance for tool selection in cfDNA tumour monitoring and establish DecoNFlow as a robust framework for benchmarking and applying DNAm deconvolution.

## Introduction

Circulating cell-free DNA (cfDNA) is DNA that is released by cells in body fluids through processes such as apoptosis, necrosis or NETosis^1^. In cancer patients, DNA derived from tumour cells is shed into body fluids where it mixes with cfDNA released by healthy, mostly immune, cells^2^. Cancer-specific DNA methylation (DNAm) markers are preserved in the tumour derived cfDNA fraction and therefore can be detected in blood plasma samples^3^, opening opportunities for several applications in cancer diagnostics and therapy monitoring: (1) methylation profiling of plasma cfDNA has shown potential for cancer screening and early tumour detection^4^; (2) we and others have shown its application for cancer classification through tissue of origin detection^5–8^; (3) estimating the fraction of tumoral cfDNA based on methylation profiling has been shown to be useful for prognostication^2,9,10^ and (4) as an alternative to mutation-based cfDNA monitoring, as it does not require prior knowledge of tumor-specific genetic mutations.

However, estimating the tumor-derived cfDNA fraction is challenging because liquid biopsy samples contain a mixture of cfDNA from both cancer and healthy cells^2^. As a possible solution, DNAm computational deconvolution algorithms can be adopted to estimate tumoral cfDNA fractions in liquid biopsy samples. The first DNAm deconvolution tool was developed in 2012 by Houseman^11^. Subsequently, in the last decade several other tools were published, recently reviewed by Ferro Dos Santos et al.^12^. These algorithms can be broadly divided into two main classes: reference-based and reference-free methods^12^. Reference-based deconvolution algorithms rely on a reference matrix, that contains cell-type-specific DNAm signatures. They estimate the relative proportion of each cell type in the bulk samples by comparing the observed methylation patterns to those in the reference matrix. Moreover, within reference-based tools, some algorithms use uniquely the DNAm percentage (beta values) at specific CpG sites or regions (e.g. meth_atlas or Houseman), while others use raw counts adjusting for different sequencing depth (e.g. CelFiE, MetDecode) or read-level information (e.g. UXM)^12^. On the contrary, reference-free methods do not require prior information and infer cell-type composition directly from the data. They are usually adopted when no reference (tumour or normal cell types) samples are available. A downside of reference-free tools is that it is challenging to assign the right cell-type entity to a specific proportion, making reference-free methods more difficult to interpret.

Over the past four years, four benchmarking studies have evaluated the performance of deconvolution tools on methylation data obtained from Illumina methylation arrays and/or sequencing-based methylation profiling for cell-type deconvolution of bulk methylome samples^13–16^. However, none of the studies focused on the task to accurately detect the presence of tumoral fractions and on the estimation of those fractions, both important metrics for the evaluation of cfDNA methylation deconvolution in tumour monitoring studies.

In this manuscript, we present DecoNFlow, a reproducible, scalable and portable Nextflow^17^ pipeline designed for cell-type deconvolution of bulk DNA methylation sequencing data. DecoNFlow allows researchers to use ten of the most used reference-based deconvolution tools (CelFiE^18^, CIBERSORT^19^, EpiDISH^20^, EpiSCORE^21^, Houseman’s CP with equality constrain^11^, Houseman’s CP without equality constrain^11^, meth_atlas^22^, MetDecode^23^, PRMeth^24^ and UXM^25^), two reference-free deconvolution tools (MeDeCom^26^ and RefFreeCellMix^27^) and three marker selection methods (DMRfinder^28^, limma^29^, wgbstools^30^), making it the most comprehensive pipeline for sequencing-based DNAm deconvolution to date. Here, we used DecoNFlow to benchmark the performance of various deconvolution tools on in silico DNA methylation mixture samples, aiming to assess their sensitivity in detecting low tumour fractions and their accuracy in estimating these fractions. This benchmarking was performed on more than 3600 samples, covering multiple factors, including tumour types, sequencing depths, marker selection methods, tumour fractions, and methylation profiling technologies.

## Results

### Benchmarking datasets generation

We benchmarked 10 reference-based and 2 reference-free DNAm deconvolution tools (summarized in Supplementary Table 1A-B) to identify the best performing tools for tumour fraction estimation. For the benchmarking study, we used sequencing-based methylation profiles from *in silico* mixture samples, i.e. mixtures of tumour and healthy control samples as a proxy for real cfDNA samples from cancer patients (Figure 1). To be as comprehensive as possible and to simulate real-world scenarios, we compared the deconvolution tools across different conditions, such as different sequencing depths, tumour types, tumoral fractions and DMR selection approaches (Figure 1). To generate *in silico* mixed samples with known tumoral fractions, we developed SillyMix, a Nextflow pipeline that uses BBMap^31^ to subsample an exact number of aligned reads directly from BAM files. Using this pipeline, we generated three different datasets:

**(i) WGBS-TT dataset:** This dataset comprises 1,640 whole-genome bisulfite sequencing (WGBS) mixed samples generated by combining methylation data from tissues of four tumour types (TT)—breast carcinoma (BRCA), bladder carcinoma (BLCA), lung adenocarcinoma (LUAD), and lung squamous cell carcinoma (LUSC)— with WGBS data from healthy plasma cfDNA. The mixtures were simulated at four sequencing depths (62M, 183M, 310M, and 620M aligned reads) and across ten tumour fractions spanning from 0.01% to 50% (Figure 1; Supplementary Table 2D). In addition, a reference dataset containing three (or four in the case of BLCA) independent reference samples for each tumour type as well as healthy cfDNA reference samples was assembled.
**(ii) RRBS-TT dataset:** This reduced representation bisulfite sequencing (RRBS) dataset includes 1,550 mixed samples generated by combining methylation data from tissues of three tumour types (TT)—lung cancer (LC), prostate cancer (PC), and colorectal cancer (CRC)— with RRBS methylation data from healthy plasma cfDNA. The mixtures were simulated at five sequencing depths (2M, 5M, 10M, 15M, and 20M aligned reads) and across the same ten tumoral fractions as the WGBS-TT dataset. In addition, a reference dataset containing three independent reference samples for each tumour type as well as five healthy cfDNA reference samples was assembled.
**(iii) RRBS-CL dataset:** This RRBS dataset consists of 440 RRBS samples derived from a neuroblastoma (NBL) tumour cell line (CL) (CLBGA) mixed with RRBS methylation data from healthy plasma cfDNA. The samples were generated at the same sequencing depths and tumoral fractions as the RRBS-TT dataset. In addition, a reference dataset containing nine other neuroblastoma cell lines as well as healthy cfDNA reference samples was assembled.

**Figure 1:**
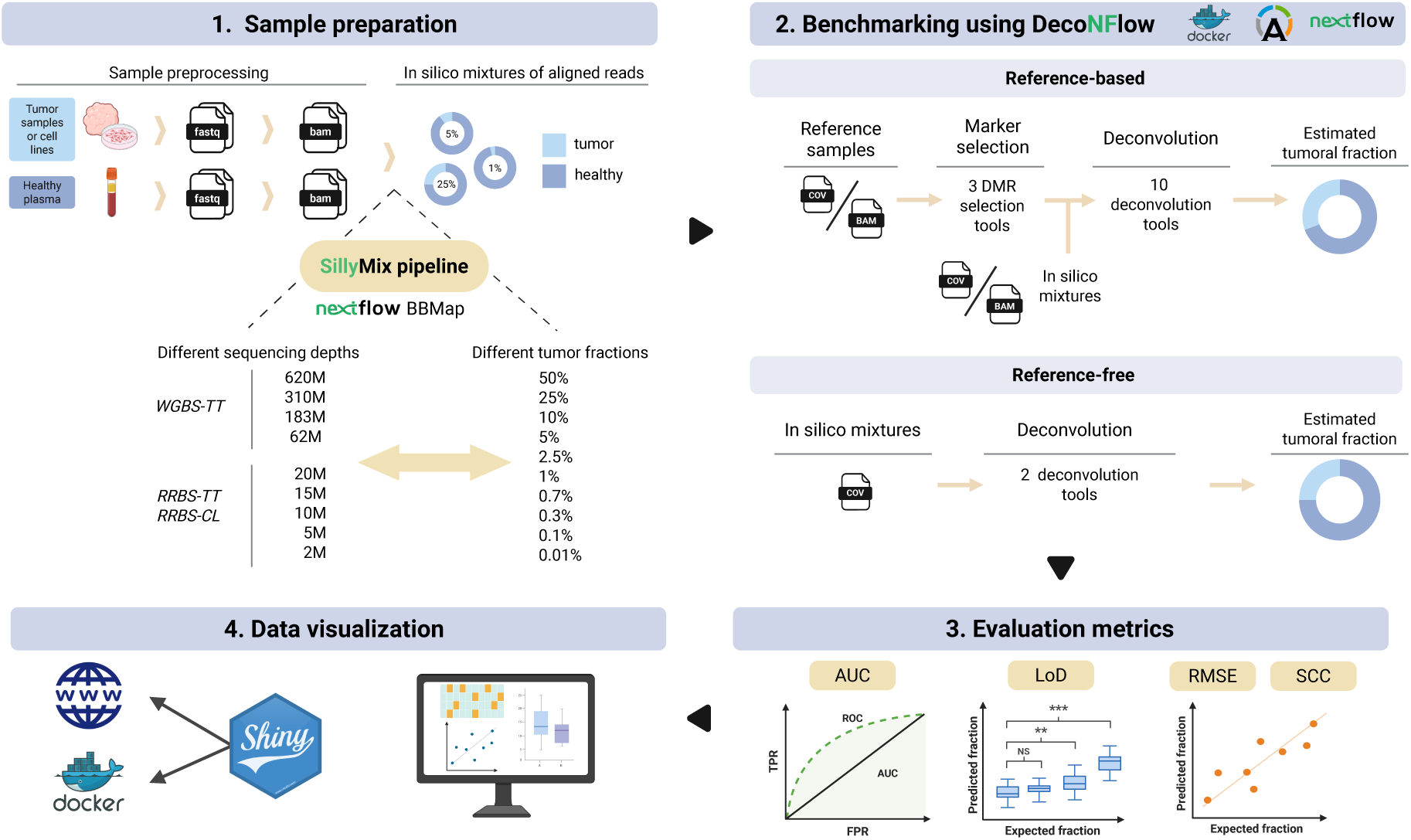
Overview of the benchmarking setup. **(1)** In this study, *in silico* mixtures of tumour tissue (TT)/cell lines (CL) samples and healthy plasma cfDNA were generated using the in-house developed SillyMix pipeline, which uses the BBMap tool to mix exact number of aligned reads from BAM files. **(2)** Then, the benchmarking was run on the mixtures using the DecoNFlow pipeline, which allowed us to estimate the tumoral fractions with 12 deconvolution tools. **(3)** The estimated fractions were compared with the expected fractions to compute four evaluation metrics (RMSE, AUC, SCC and LoD) **(4)** The results are visualized in interactive mode using an R Shiny app, which can be used through a web server or locally using a Docker container. Created with BioRender.com.

### Benchmarking set-up and execution

To run all the deconvolution tools, we developed DecoNFlow, a Nextflow pipeline where each tool evaluated in this study has been containerized. This allows the users to simultaneously deconvolve bulk samples using several DMR selection approaches and deconvolution tools in parallel, without the need of switching platforms and/or programming languages (Figure 1). Since the reference-based methods rely on reference matrices composed of tumour and healthy specific DMRs, we evaluated deconvolution performance across three different DMR selection approaches (Supplementary Table 1C). We constrained each DMR selection tool to select the top 250 hypo-methylated regions per entity with adjusted p-value < 0.01 (Materials and Methods). On the other hand, since reference-free tools directly model the bulk samples to perform deconvolution without a reference matrix, we evaluated the performance of these tools across two different approaches to aggregate the individual CpGs into regions (Materials and Methods).

To evaluate the performance of the different tools, estimated and expected tumour fractions are compared using four different metrics: (i) the area under the Receiver Operating Characteristic curve (*AUC) value* to measure the capability of a tool to predict presence or absence of tumoral signal; (ii) the limit of detection (LoD) to measure the lowest tumoral fraction the deconvolution tools can still detect; (iii) root-mean squared error (*RMSE)* to measure the accuracy of the estimated tumoral fractions; and (iv) Spearman’s rank correlation coefficient (*SCC)* to measure the correlation between the estimated and the expected tumoral fractions.

Finally, to allow researchers to easily explore all the benchmarking results in more details and in an interactive way, we developed an R Shiny app^32^.

### Overall performance of reference-based deconvolution tools

To compare the performance of each tool, we performed an overall ranking of the different deconvolution methods across the different datasets (different sequencing libraries, sequencing depths and different tumour types). Briefly, for each dataset analysed, the metrics (SCC, 1-RMSE, AUC and LoD) were min-max scaled across deconvolution tools at each combination of sequencing depth, tumour type and DMR selection approach adopted. Then, for each deconvolution tool, the scaled values were averaged across all combinations to obtain an overall score per dataset. Finally, to get a final overall performance score per deconvolution tool, a weighted mean of each dataset’s score was computed (Materials and Methods). This analysis revealed that CelFiE is the best overall performing tool, robustly ranking in the first position across different conditions and scenarios (Figure 2). The two reference-free methods, RefFreeCellMix and MeDeCom, rank second and third, respectively. However, it is important to note that while these two tools have the best SCC, AUC and LoD scores, they are the worst performers in terms of RMSE, due to their consistent over-estimation of tumoral fractions. Houseman’s CP with and without equality constrain (Houseman_eq and Houseman_ineq), meth_atlas and MetDecode also show very good performance across the different DMR selection tools and datasets, composed of samples generated using different sequencing approaches (Figure 2, Supplementary Figure 1-6). Notably, while UXM ranks among the worst tools in the RRBS-TT and RRBS-CL datasets, it is among the best methods on WGBS-TT dataset (Figure 2, Supplementary Figure 1-6).

**Figure 2.**
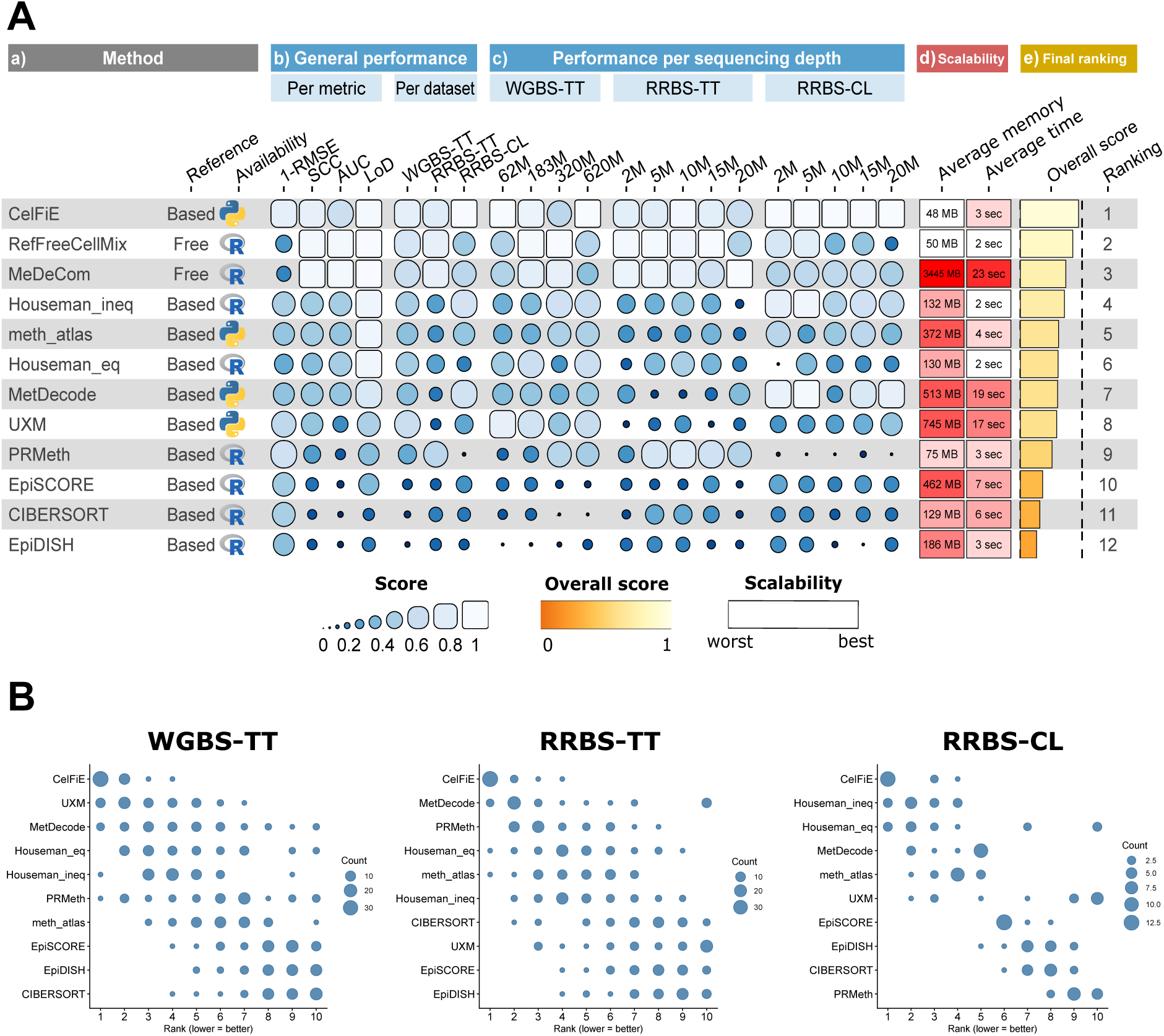
Summary of the benchmarking results of the reference-based and -free deconvolution tools. **(A)** Funkyheatmap showing the deconvolution benchmarking results. This heatmap includes: (a) the deconvolution tools ordered by overall score, the type of tools (reference based vs free) and the programming language originally used to develop the tool; (b) the performance of the tools aggregated by metric and by dataset, respectively; (c) the performance aggregated by sequencing depth for each dataset; (d) the scalability in terms of memory and run time and (e) the final ranking based on the overall score computed as the weighted mean of individual datasets’ overall scores and scalability. Aggregated scores are min-max scaled per column, only enabling comparison of each tool’s performance within one tested condition. **(B)** The rank distributions of the overall score of reference-based tools, shown separately for each dataset, highlight how tool performance varies across datasets. To generate the rank distributions, the overall score was computed at each combination (DMR tool, sequencing depth and tumour type) and the tools ranked from highest to lowest score. Finally, for each deconvolution tool the rank distribution was plotted, where the dot size indicates how often a tool was assigned to a given rank.

In the next sections, we analyse the results for each metric separately to better highlight the specific strengths and limitations of each tool. However, since reference-free and reference-based methods require a different set-up, we first analyse the performance of the reference-based methods and finally compare them with the reference-free methods in a separate section.

### Comparison of the accuracy in predicting presence/absence of tumoral fraction using the AUC metric

An important challenge of computational deconvolution of cfDNA methylomics is to accurately predict the presence or absence of tumoral-derived DNA in a bulk cfDNA sample. To quantify this, we measured the AUC value in a ROC analysis for different sequencing depths and tumoral fractions of the samples, as AUC is expected to be impacted by those parameters. Based on the AUC value across all conditions, CelFiE is consistently the best reference-based tool (Figure 2-3, Supplementary Figure 7). As expected, increasing the sequencing depth in WGBS-TT consistently improves the AUC values in the case of CelFiE, MetDecode, Houseman_eq, Houseman_ineq, UXM and PRMeth (Figure 3A, Supplementary Figure 8). However, when looking at the RRBS-TT dataset, increasing sequencing depth beyond 10M reads does not yield a significant improvement in AUC (Figure 3B, Supplementary Figure 9). In RRBS-CL dataset this effect is even stronger, where apart from UXM no significant improvement in the AUC value is observed above a sequencing depth of 5M reads (Figure 3C). We speculate that this is due to sequencing saturation, which we define as the point from where increasing sequencing depth will mostly lead to an increase in (PCR) duplicated reads. Since in RRBS it is not possible to deduplicate the samples during the preprocessing, beyond 10M reads, PCR duplicates are assumed to accumulate, increasing noise, and the AUC value does not improve anymore. This hypothesis is supported by the sequencing saturation plots shown in Supplementary Figure 10, which clearly demonstrate that, within the tested range of sequencing depths, the RRBS datasets have reached saturation, whereas the WGBS dataset has not. Achieving saturation for the latter would require substantially higher sequencing depths, reaching levels that are costly prohibitive.

**Figure 3.**
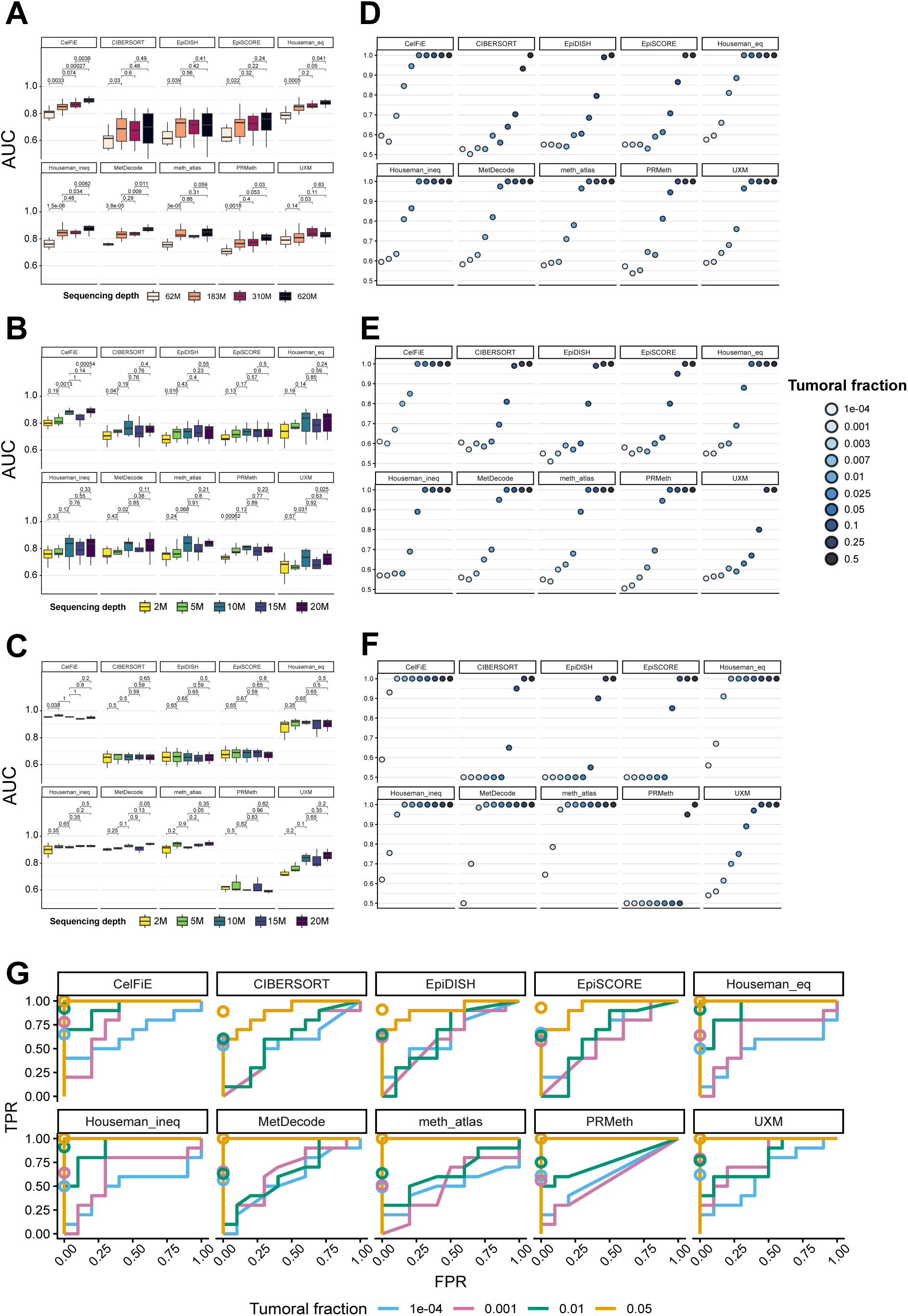
Predicting presence of tumoral fraction: The raw AUC scores for the **(A)** WGBS-TT, **(B)** RRBS-TT and **(C)** RRBS-CL datasets. On the left side, boxplots show the AUC score distribution per tool for each sequencing depth. Unpaired Mann-Whitney U test was used, adopting Benjamini-Hochberg correction for multiple testing, to test whether the AUC score is significantly different between the sequencing depths. The median AUC score at each tumoral fraction was plotted for WGBS-TT **(D)**, RRBS-TT **(E)** and RRBS-CL **(F)**. **(G)** An example of ROC analysis for 4 low fractions of the mixed samples of LUAD tumour data in a non-tumoral background (620M sequencing depth, limma). The circle dots indicate the AUC value for each tumour fraction (visit the R Shiny app for more interactive visualizations).

Next, we compared the AUC values across different tumoral fractions. For all the WGBS-TT and RRBS-TT datasets, all the tools have a median AUC lower or equal to 0.6 at the lowest fraction (1e-04), reflecting an almost random classification (Figure 3D and E). At 0.05 tumoral fraction, the top performing methods (CelFiE, Houseman_eq, Houseman_ineq, meth_atlas, MetDecode and UXM) reach a median AUC equal to 1, showing a perfect discrimination between the tumour and healthy samples (Figure 3D and E). One exception is UXM, where for the RRBS-TT dataset an AUC of 1 is only reached at 0.1 tumoral fraction (Figure 3E). For the RRBS-CL dataset, this is more outspoken with CelFiE reaching an AUC of 1 at 0.003 already (Figure 3F), the other top performing tools (Houseman_eq, Houseman_ineq, meth_atlas and MetDecode) at 0.007 and UXM only at 0.1.

### The tumoral fraction limit of detection

In the context of tumour monitoring applications, it is important to know the lowest tumoral fraction that deconvolution tools can detect, i.e. limit of detection (LoD). To this end, we performed, for each tumoral fraction, a one tail Mann-Whitney U test (adjusted p-value < 0.01, Benjamini–Hochberg (BH) correction) to identify the lowest tumour fraction where the estimated fraction distribution (n = 10 replicates) is significantly higher than the estimated fraction distribution on control samples (0% tumour fraction) (Figure 4B and C). We also studied LoD in relation to sequencing depth.

**Figure 4.**
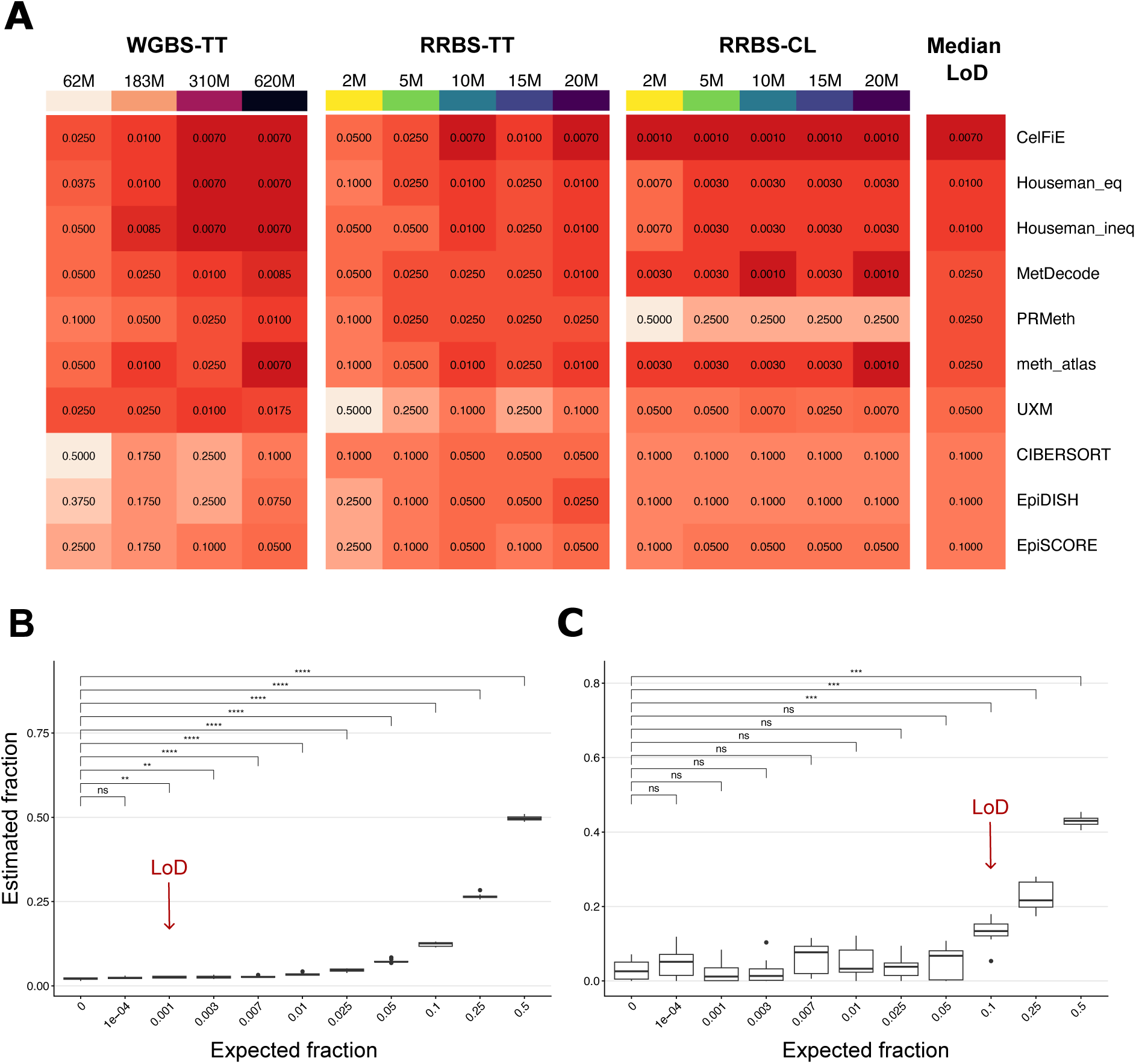
Lower limit of detection (LoD) of tumour fractions: **(A)** The heatmaps show the median LoD of the tools for different sequencing depths of each dataset (WGBS-TT, RRBS-TT and RRBS-CL). The deconvolution tools are ranked by increasing overall median LoD (last column) which is the overall median LoD of the three datasets. **(B and C)** The LoD is computed as the lowest tumour fraction where the estimated fraction distribution of the 10 replicates is significantly different from the estimated fraction distribution in the samples with 0% tumour fraction (adjusted p-value < 0.01, Unpaired Mann-Whitney U-test with BH correction). Here the LoD is calculated for the LUAD tumour type (WGBS-TT dataset), with 620M sequencing depth, using wgbstools as DMR tool for CelFiE **(B)** and EpiDISH **(C).**

Also for this metric, CelFiE is the top performing reference-based tool, showing the lowest median LoD in the RRBS-TT (0.7% at 10M reads), in WGBS-TT together with Houseman_eq and Houseman_ineq (0.7% at 310M reads) and in RRBS-CL (0.1% at 2M reads) datasets (Figure 4, Supplementary Table 3A). Notably, in WGBS-TT, almost for all the tools the median LoD keeps on improving (i.e. lowering) with increasing sequencing depth, except for the best performing tools, i.e. CelFiE, Houseman_eq and Houseman_ineq, where the minimal LoD (0.7%) is reached at 310M sequencing depth (Figure 4A, Supplementary Table 3A). In contrast, LoD already reaches a plateau at 10M reads sequencing depth for most tools in the RRBS-TT dataset, apart from PRMeth and MetDecode, whose lowest LoD is reached at 5M and 20M, respectively. Similarly, in RRBS-CL dataset the LoDs plateau is reached at 5M or 10M for all the tools, apart from MetDecode that reaches it at 20M. These results align with the AUC results, suggesting again that these differences can be explained by the fact that the sequencing saturation is not reached for the WGBS dataset compared to the RRBS datasets.

### Accuracy in tumour fraction prediction (1-RMSE and SCC)

To allow precise monitoring of tumoral cfDNA fractions, an accurate estimation of the tumoral fractions through deconvolution is necessary. To evaluate this performance parameter, we used both the 1-RMSE and SCC metrics.

Since RMSE is not a scale-free metric, samples with higher tumour fractions will show higher RMSE values and will therefore contribute more heavily to the overall RMSE. Consequently, the overall RMSE tends to be skewed towards the results of the higher tumour fractions, as illustrated in Supplementary Figures 2, 4, and 6. To mitigate this bias, each dataset was stratified into three tumour fraction classes: low (≤2.5%), medium (2.5% to 25%), and high (≥25%). The RMSE was then computed within each of these ranges for every combination of tumour type, sequencing depth and DMR tool.

Overall, CelFiE showed the highest median 1-RMSE at all tumour fraction classes (Figure 5A) and the best 1-RMSE rank distribution (Figure 5C). For the WGBS-TT dataset, CelFiE shows the best performance at both medium and high fractions (Supplementary Figure 11), however PRMeth ranks first for the low fractions, followed by UXM and CelFiE. In the RRBS-TT, CelFiE performs best with respect to 1-RMSE across all tumour fractions (Supplementary Figure 11), while in the RRBS-CL dataset CelFiE, Houseman_eq, Houseman_ineq and meth_atlas performs best with high 1-RMSE values for both low and medium fractions, but EpiSCORE and meth_atlas performs best at high fractions (Supplementary Figure 11).

**Figure 5.**
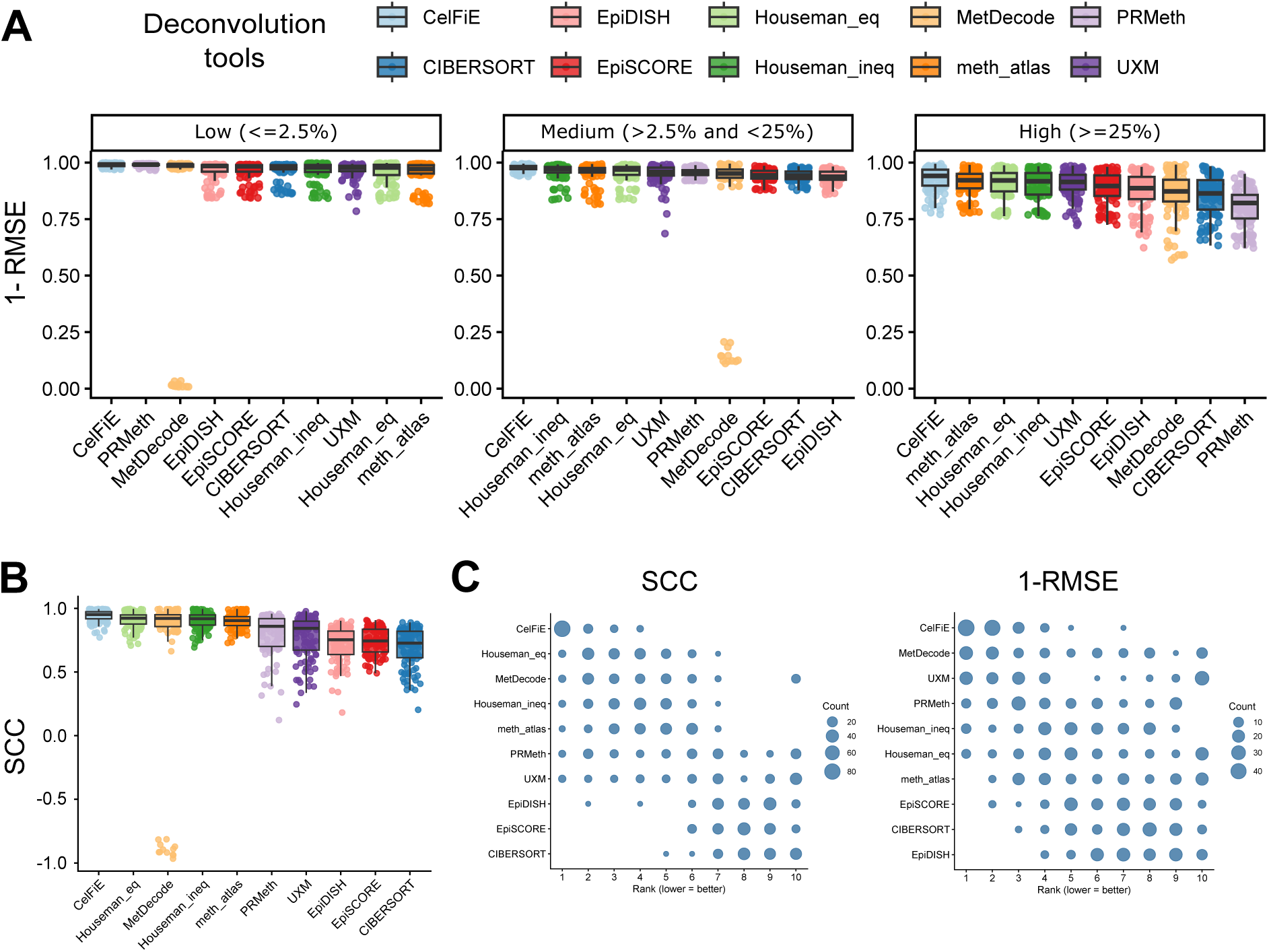
Accuracy of tumoral fraction estimations by deconvolution tools evaluated with raw 1-RMSE and SCC on combined WGBS-TT, RRBS-TT and RRBS-CL datasets. **(A)** For 1-RMSE analysis, the data was split into three tumour fraction classes: low (≤2.5%), medium (>2.5% and <25%), and high (≥25%). The raw 1-RMSE was computed at each combination of tumour type, sequencing depth and DMR tool and plotted in boxplots, finally ordered by decreasing median 1-RMSE. **(B)** Boxplots representing the raw SCC, computed across all tumoral fractions, ordered by decreasing median SCC. **(C)** The rank distributions of SCC and 1-RMSE, shown over all datasets, were calculated in the same way as described in Figure 2B. The dot size indicates how often a tool was assigned to a given rank.

In addition, SCC was calculated across all tumour fractions. Over all datasets, CelFiE consistently achieved the highest median SCC (Figure 5B) and SCC rank distribution (Figure 5C), with Houseman_eq, Houseman_ineq, meth_atlas, MetDecode and UXM on the remaining top 5 positions. Noticebly, MetDecode shows anticorrelated predictions for a small number of outlier samples (Figure 5, Supplementary Figure 11).

With respect to sequencing depth, SCC significantly increases at higher depth, consistent with the trends observed for AUC and LoD. A similar upward trend is seen for 1-RMSE; however, this change is not statistically significant, likely because most of the expected tumour fractions are very low, resulting in only minimal differences (Supplementary Figure 12).

### Reference-free vs reference-based tools

In this benchmarking study we also included two reference-free deconvolution tools: MeDeCom and RefFreeCellMix. These methods are specifically of interest in case reference methylation data are lacking for certain tumour or tissue types.

MeDeCom and RefFreeCellMix outperform all the reference-based deconvolution tools in terms of raw AUC value, LoD and SCC when looking at the overall scores (Figure 6A). More specifically, this is also the case when we look at the results for the WGBS-TT and RRBS-TT datasets separately (Supplementary Figure 13A-B), while for the RRBS-CL dataset CelFiE ranks first (Supplementary Figure 13C). In contrast, when looking at the raw 1-RMSE, both MeDeCom and RefFreeCellMix underperform compared to all the reference-based tools, even the worst performing ones (Figure 6B). These results are explained by a consistent over-estimation of the tumoral fraction by both MeDeCom and RefFreeCellMix (Figure 6C, Supplementary Figure 13A-C). These results align with previous benchmarking results and with the performance evaluation of MeDeCom when applied to deconvolve 2 latent components/entities^14,26^. Lastly, while these tools applied on the RRBS-TT and RRBS-CL datasets show similar metrics compared to reference-based tools, their AUC and SCC on the WGSB-TT dataset plateau at 310M reads (Supplementary Figure 14). This suggests that for WGSB data, a sequencing depth of 310M may be sufficient to achieve optimal performance for reference-free tools (Supplementary Figure 14). These findings together indicate that, while MeDeCom and RefFreeCellMix can capture tumour signal sensitively and robustly (high AUC, high SCC, low LoDs), they are poorly calibrated in absolute terms when applied to deconvolve bulk samples into 2 entity proportions, showing very low accuracy (1-RMSE), and highlighting the importance of a reference for precise tumour fraction estimation.

**Figure 6.**
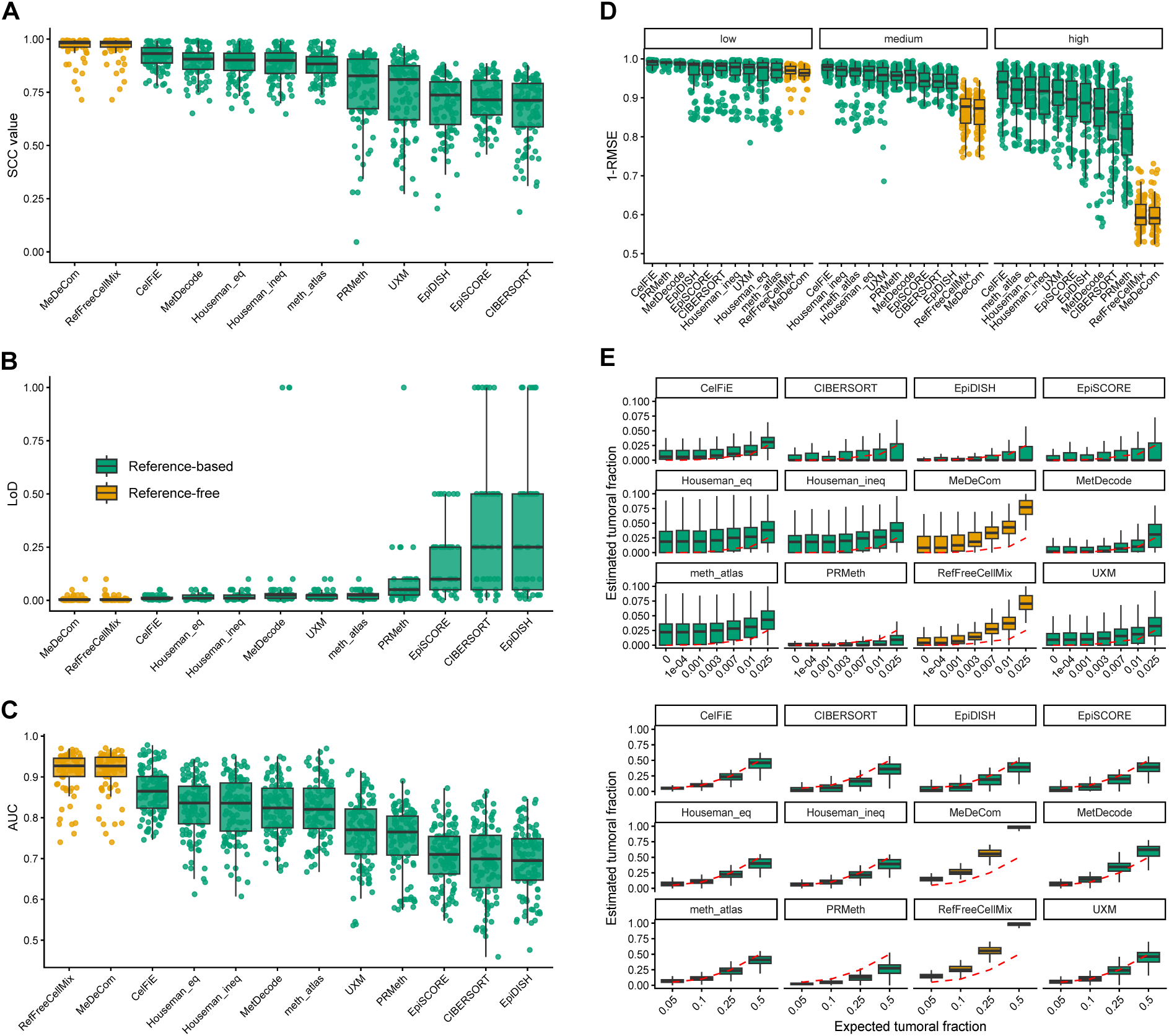
**(A,B,C)** Boxplots showing, respectively, the SCC, LoD and AUC value distributions for all combinations and datasets across the different deconvolution tools. **(D)** 1-RMSE values summarized for all combinations and datasets across the different tumour fraction classes, i.e. low (<=0.025), medium (>0.025 and <0.25) and high (>=0.25). **(E)** Boxplots per deconvolution tool showing estimated vs expected tumoral fractions divided into two plots to better visualize low (<=0.025, up) and high (>0.025, bottom) fractions, and the consistent over-estimation of tumoral fractions from MeDeCom and RefFreeCellMix.

### Computational resources and scalability

Next to the performance of the deconvolution tool, the computational resources needed to perform the deconvolution, such as memory peak and runtime, are important parameters to consider upon method selection. In this analysis we only focused on the computational resources needed for deconvolution and not to preprocess the data. We selected the RRBS-CL dataset and ran deconvolution on different combinations: 5, 20, 50 or 100 mixed samples; and using data from 25, 50, 100, 250 or 500 DMRs (or regions for reference-free tools) per entity to assess how well the tools scale. We allocated 8 CPUs and 16GB of memory to all the tools, which are the minimum resource requirements to run the pipeline and which are usually available in a personal laptop. All the tools, apart from MeDeCom, have a very small runtime and memory usage as averaged over all the combinations, which is expected since they are all resource-efficient models. Notably, UXM has a linear increase of runtime when increasing the number of samples, probably related to its *deconv* function, which uses bulk *PAT files* instead of *COV files* and performs deconvolution sample by sample (Figure 7C-D). Peak memory used by the tools is always below 1.5 GB, apart from MeDeCom where memory usage linearly increases at increasing number of regions and number of samples (Figure 7A-B). Overall, UXM, MeDeCom and MetDecode are among the tools that require the most computational resources, while CelFiE ranks best in terms of memory and RefFreeCellMix ranks the best in terms of running time (Figure 7A-C).

**Figure 7.**
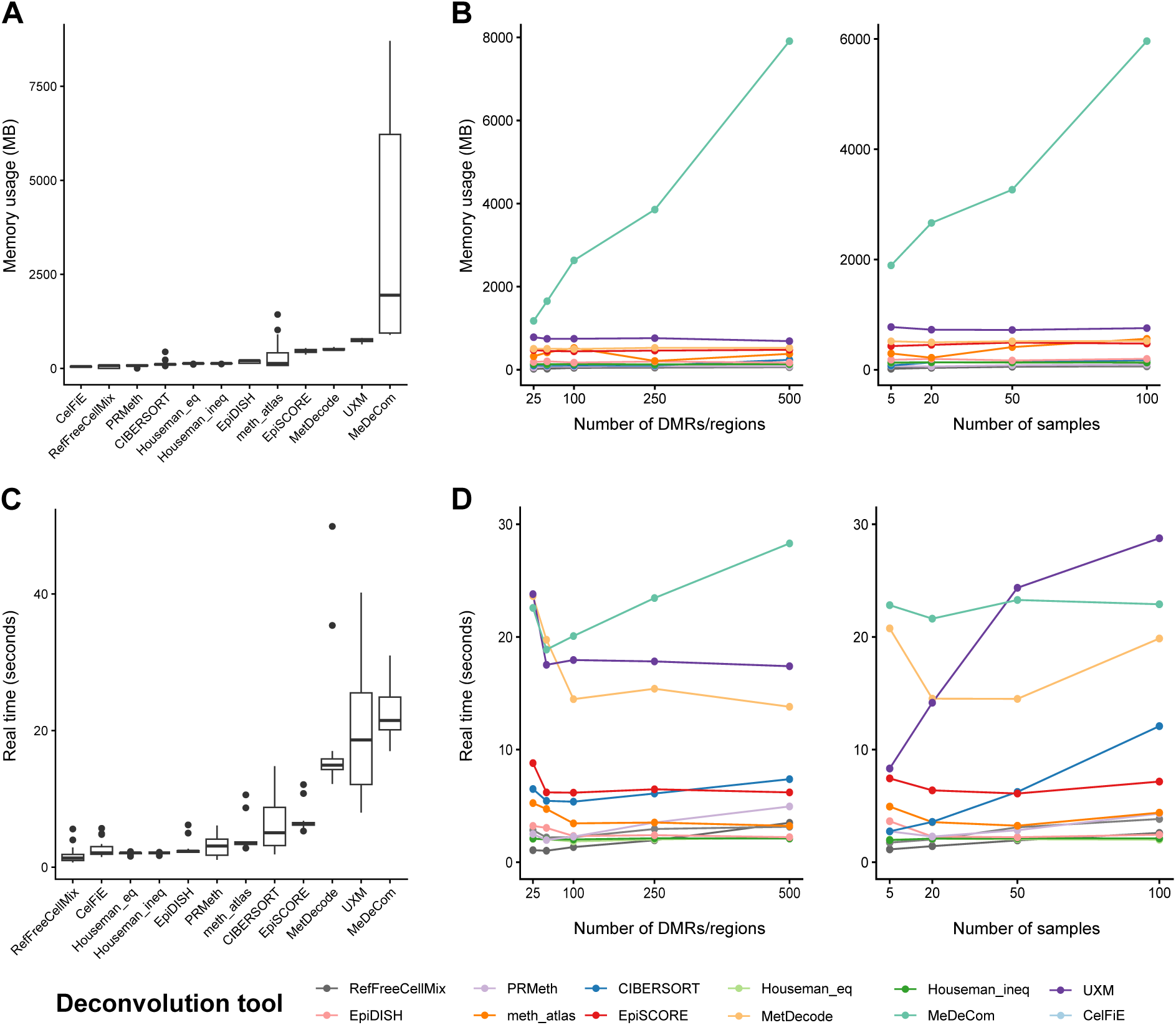
Scalability of the deconvolution tools: running time and memory usage. Each tool was executed with fixed computational resources (8 CPUs and 16 GB RAM) on subsets of the RRBS-CL dataset, varying the number of DMRs/regions per entity and the number of samples. Panels **(A) and (C)** show the overall distributions of memory usage and runtime, respectively, using boxplots, while panels **(B) and (D)** display the corresponding mean values across the tested conditions.

## Discussion

In this study, we performed a comprehensive benchmark of ten reference-based and two reference-free DNA methylation deconvolution tools for tumoral fraction estimation in samples with a healthy cfDNA background under different scenarios.

To efficiently perform this benchmarking study, we have built DecoNFlow, a Nextflow pipeline allowing to reproducibly and reliably run and evaluate deconvolution tools. This pipeline incorporates ten reference-based and two reference-free deconvolution tools. Moreover, it allows users to run an entire deconvolution workflow, since it also includes three different DMR selection methods to build the reference matrix needed for reference-based deconvolution.

To perform this benchmarking, we generated 3690 in silico mixtures of tumoral cfDNA methylation reads and healthy donor cfDNA methylation reads divided into three different datasets. These datasets were composed of data from different tumour types, two bisulfite sequencing approaches (WGBS and RRBS) and two types of starting material (tumour tissue (TT) and tumour cell line (CL)): WGBS-TT (BRCA, BLCA, LUAD and LUSC), RRBS-TT (CRC, LC and PC) and RRBS-CL (NBL).

Overall, the benchmarking study revealed that CelFiE is the best performing tool for tumoral fraction estimation across all the datasets used, across the three DMR selection methods used and across different simulated sequencing depths. Additionally, meth_atlas, MetDecode, Houseman_eq and Houseman_ineq show a very good performance, while CIBERSORT, EpiDISH, EpiSCORE and PRMeth show the worst performances overall. Regarding UXM, it performs as good as CelFiE in WGBS-TT dataset, while it underperforms in both RRBS datasets. This observation might be explained by the fact that UXM was developed and applied on deconvolution of WGBS samples in the original paper^25^, thus internal parameters might have been optimized for this data type. These results highlight the importance of diversifying the datasets, as done in this study, and shows that tool selection should be done according to sequencing technology.

Surprisingly, the two reference-free methods, MeDeCom and RefFreeCellMix outperformed all reference-based methods in terms of detection of the presence of tumour (AUC and LoD) and correlation between the estimated and actual fractions (SCC). An exception is CelFiE outperforming both reference-free methods on the RRBS-CL dataset. This observation could be attributed to the cell line based reference used for the RRBS-CL dataset, implying an exclusively tumoral composition compared to tumor tissue samples that also contain non-tumoral cells. The reference for RRBS-CL is thus inherently less heterogenous, leading to a better performance of reference-based methods, like CelFiE. While outperforming with respect to AUC and LoD, one important downside of the reference free methods is that they both show a consistent over-estimation of tumoral fraction, making them the overall worst tools in terms of 1-RMSE. The results obtained on the 1-RMSE are in line with previous benchmarking of reference-free methods when used to deconvolve bulk samples into two cell-type/entity proportions and with the results showed by the authors of MeDeCom^14,26^. Moreover, another downside is the less straightforward annotation of the deconvolved fractions to the corresponding cell-type/entity, especially when no prior information is available. These results could justify the use of one of the two reference-free methods to sensitively detect whether there is a detectable tumoral fraction, and -in a second stage- when a tumour signal is detected, use CelFiE to accurately estimate the fraction.

Second, in this study we have also shown that an important parameter impacting the overall performance of deconvolution is the sequencing depth. We found that all metrics used in this study are affected by sequencing depth. The performance improvement with increasing sequencing depth was only observed in case the sequencing saturation depth was not reached yet. For WGBS-TT, a constant increase of the metrics was observed with increasing sequencing depth, confirmed by the sequencing saturation plot that did not reach the plateau phase yet. However, when looking at the RRBS datasets, both metrics did not improve anymore with sequencing depths higher than 10M reads. In fact, in these datasets, when looking at sequencing saturation plots, at 10M reads the curves already reached the plateau phase. This finding shows the importance of sequencing saturation estimation when using RRBS data for tumour fraction estimation, in order to avoid unnecessary sequencing costs.

Given the fact that no grand differences could be observed for computational resources needed to run the different deconvolution tools, this factor should not be decisive for tool selection. Even though UXM and MetDecode demand higher computational resources, their requirements are still within a feasible range, making them practical to run in typical computing environments. An exception is the reference-free tool MeDeCom, which has a strong linear increase of memory usage at increasing number of regions and samples.

An additional aspect to highlight in this benchmarking study is the *in silico* mixture generation, which was not always reported clearly in previous studies but important as we explain below. Two main approaches exist to make in silico mixes with known tumoral fractions from pure samples, i.e; pre- and post-mapping mixing. In the first approach, reads are mixed at the FASTQ level (e.g. using seqtk^33^), which best mimics real data^34^. However, it can introduce bias due to variable mapping rates between samples generated in different batches. This can potentially lead to over- or under-representation of sequencing reads of certain samples, undermining the expected ground truth. In the second approach, pure samples are first preprocessed, and the aligned (and deduplicated, for WGBS) reads are then mixed at the BAM level^34^. This approach, on the other hand avoids mapping-related biases. Tools like SAMtools can subsample BAM files efficiently, but it do not allow to subsample an exact number of reads, as users can only specify the probability of keeping each read. Ultimately, this will not result in the expected ground truth, an aspect which is particularly problematic when mixing low fractions. For this reason, we developed SillyMix, a Nextflow pipeline which uses BBMap’s *reformat.sh* function. This BBMap function allows precise subsampling by specifying the exact number of reads, enabling generation of in silico samples with more accurate ground truth.

Although this study provides important guidelines for DNAm computational deconvolution, some limitations should be considered. First, this benchmarking has been conducted exclusively on *in silico* mixtures. While this approach allows us to derive a ground truth and to precisely evaluate the deconvolution tool performances, it may also oversimplify the biological complexity of real mixtures, potentially leading to a general overestimation of accuracy. Nonetheless, assessing tumour fraction estimation on actual plasma cfDNA samples remains highly challenging given the difficulty to know the ground truth. Second, this study relied exclusively on bisulfite-converted methylome samples. We do not foresee differences using enzymatic conversion approaches, however additional methylomics approaches such as Oxford Nanopore Technologies^35^ needs to be incorporated in the future when long-read sequencing of cfDNA becomes more routinely implemented. However, nowadays, tumour datasets profiled with this technology are still limited. Third, for the WGBS-TT dataset, *in silico* mixtures were not generated at sequencing depths sufficient to achieve saturation. Although this prevented assessment of tool performance at near-saturation levels, in most liquid biopsy studies sequencing saturation is rarely reached due to practical and resource limitations, particularly in a clinical setting^36–38^. Finally, rather than identifying one best DMR selection tool, we evaluated the deconvolution methods across diverse scenarios to provide a broader assessment of their performance.

In summary, based on the datasets we analysed, we advise to use CelFiE for tumoral fraction estimation in case a reference dataset is available. Ideally, this reference dataset contains data from a pure population of tumoral or healthy cells. Our benchmarking results also point at the value of reference-free methods, suggesting the potential benefit to complement the results of reference-based methods with those of reference-free methods, to aid in the estimation of the presence of a tumoral signal in a sample. However, to estimate the fractions of the tumoral DNA in the mixed sample, the use of reference-free methods is highly discouraged, since these methods show a consistent over-estimation of the fractions. Within this study, we also present DecoNFlow, a Nextflow pipeline that allows to run and compare multiple deconvolution tools in parallel on bulk samples, additionally allowing to integrate each estimation to an ensemble result of multiple tools. DecoNFlow has been used here to perform a benchmarking of deconvolution tools for tumor fraction estimation. However, it can also be used for general cell-type deconvolution of bulk methylome samples, providing a unified and containerized environment for reproducible experiments. Finally, in our study we also implemented an R Shiny app that researchers can consult to look into more detail in the results, complementing the information and guidelines we have provided here, further aiding them to make a well-thought deconvolution tool selection.

## Materials and Methods

### Datasets

#### Whole-genome bisulfite sequencing tumour tissues dataset (WGBS-TT)

##### Human primary tumour samples

To build the whole-genome bisulfite sequencing tumour tissue (WGBS-TT) dataset we used data from 21 samples from 4 different tumour types: breast carcinoma (BRCA, n=5), bladder carcinoma (BLCA, n=6), lung adenocarcinoma (LUAD, n=5) and lung squamous cell carcinoma (LUSC, n=5). Data was downloaded from TCGA using the corresponding manifests present at the GDC Data Release 36.0 (https://docs.gdc.cancer.gov/Data/Release_Notes/Data_Release_Notes/#data-release-360), controlled and maintained by dbGaP^39^ (Supplementary Table 2A). Samples were available in BAM format. To convert them to a FASTQ format, the function *bamtofastq* from BEDtools^40^ (v2.31.0) was used. WGBS data from three (BRCA, LUAD and LUSC) or four (BLCA) tumour samples was used for the reference matrix construction (reference set). The remaining two tumour samples were merged into a unique FASTQ file used to generate in silico bulk mixtures (testing set, Supplementary Table 2A).

##### Healthy plasma cell-free DNA

Healthy plasma cell-free DNA from 12 healthy donors profiled with WGBS were downloaded from SRA Run Selector (https://www.ncbi.nlm.nih.gov/Traces/study/?acc=PRJNA691320&o=acc_s%3Aa)^18^ (Supplementary Table 2A). WGBS from 5 healthy samples were used for reference matrix construction (reference set), while the remaining 7 healthy samples were merged into a unique FASTQ file to generate in silico bulk mixtures (testing set, Supplementary Table 2A).

##### Sample preprocessing

First, and only for the reference set, to reduce potential sequencing bias, both the tumoral and healthy FASTQ files with more than 315M of paired-end reads were subsampled to 315M. Second, both the reference and testing set FASTQ files were trimmed, aligned (to the human genome version hg19), deduplicated and the methylation status was extracted using nf-core/methylseq pipeline (v.2.6.0)^41,42^ and using the (default) Bismark^43^ subworkflow. For the healthy samples, we used the following additional parameters for preprocessing, as described in the original publication^18^: *--clip_r1 4 --clip_r2 4 --three_prime_clip_r1 12 --three_prime_clip_r2 12.* Original query sorted BAM files were coordinated sorted and indexed using SAMtools^44^ (v1.19.2).

#### Reduced-representation bisulfite sequencing datasets (RRBS-TT and RRBS-CL)

##### Human primary tumour samples and cell lines

Reduced-representation bisulfite sequencing (RRBS) data were collected from both primary tumour tissues and neuroblastoma cell lines.

The RRBS tumour tissue dataset (RRBS-TT) included 15 samples from three tumour types: lung cancer (LC, n=5), colorectal cancer (CRC, n=5), and pancreatic cancer (PC, n=5), downloaded from SRA Run Selector (PRJNA315188)^45^ (Supplementary Table 2B). RRBS data from three tumour samples was used for the reference matrix, while the remaining two tumour samples were pooled into a single FASTQ file to generate the *in silico* mixtures (Supplementary Table 2B).

The RRBS cell line dataset (RRBS-CL) comprised nine neuroblastoma (NBL) cell lines which were profiled using cfRRBS and used to construct the reference matrix (Supplementary Methods). One additional cell line (CLB-GA) was cultured and used to generate artificial cfDNA (arti-cfDNA) through MNase digestion (Supplementary Methods). Next, arti-cfDNA was used to perform cfRRBS (Supplementary Methods), and the resulting DNA methylation profile were used for the *in silico* mixture generation (Supplementary Table 2C).

##### Healthy plasma cell-free DNA

A total of 11 healthy cfDNA donor samples was downloaded from EGA^46^ (EGAD50000000734^5^) and used to create the reference dataset (Supplementary Table 2B). 15 additional plasma cfDNA samples were processed in-house using cfRRBS (see Supplementary Methods), pooled together into a single composite sample and then used to generate the *in silico* mixtures (Supplementary Table 2B-C).

##### Samples preprocessing

All RRBS datasets (tumour tissues, cell lines, and healthy cfDNA FASTQ files) were trimmed, aligned (human genome version hg19), optical duplicates were removed and methylation status extracted using an in-house developed Nextflow pipeline. In brief, nf-core/methylseq pipeline was slightly modified to add a module in the Bismark workflow for the removal of optical duplicates using Picard Tools^47^. We used the version v2.6.2 of the pipeline, which can be found at https://github.com/VIBTOBIlab/MethylSeq with the following additional parameters: *--rrbs true --clip_r1 3 --clip_r2 4 --three_prime_clip_r1 1 --three_prime_clip_r2 1.* Original query sorted BAM files were coordinated sorted and indexed using SAMtools^44^ (v1.19.2).

#### Generation of the *in silico* mixtures

*In silico* mixtures were generated following a consistent procedure across all datasets. Preprocessed BAM files, obtained from the merged FASTQ files retained for *in silico* mixture generation, were processed with the in-house developed Nextflow SillyMix pipeline (v1.1.0, https://github.com/VIBTOBIlab/Sillymix), which utilizes the BBMap software^31^ to specify the exact number of reads to subsample from a BAM file (see Discussion for more details).

The healthy and tumour samples were mixed at 10 different tumour fractions and 4 different sequencing depths (Supplementary Table 2D). For each tumour fraction, 10 replicates were generated, with sampling performed using a different seed each time for both the healthy and tumour samples. Since the number of base-pairs influences the actual tumoral fraction, we corrected the number of reads subsampled for the average read length of the samples that were being mixed. To correct for this bias, we aimed to solve the following system:

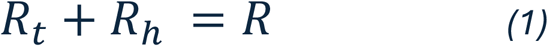

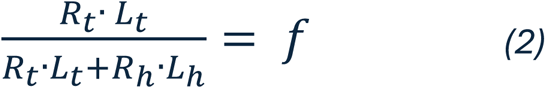

Where *R* is the total number of reads to subsample (sequencing depth), *f* is the desired tumour fraction in base pairs, *R*_*t*_ *and R*_*h*_ is the number of tumour and healthy reads to sample and *L*_*t*_ *and L*_*h*_ is the average read length of the tumour and healthy samples, respectively. Since *L*_*t*_, *L*_*h*_ *and R* are known, solving the system finds the number of tumour and healthy reads corrected for the average read length. Finally, we used Bismark with the *--single parameter* to extract methylation values, as the paired-end information is lost during subsampling with BBMap.

In total, 400 *in silico* mixtures were generated per tumour type in WGBS-TT and 500 in RRBS-TT/RRBS-CL, respectively (10 replicates * 10 tumour fractions * 4 or 5 sequencing depths, respectively), summing up to 1600 *in silico* WGBS-TT mixtures, 1500 *in silico* RRBS-TT mixtures and 500 RRBS-CL *in silico* mixtures. Additionally, 10 healthy replicates (0% tumour fraction) were generated per sequencing depth: in total, 40 for WGBS-TT and 50 for RRBS-TT and RRBS-CL. In total, 3690 in silico mixtures were generate and used for the benchmarking.

### Generation of a reduced-representation human genome (hg19)

To convert the hg19 reference genome to a reduced-representation format (hg19-RR), we used <u>mkrrgenome</u>. After initial conversion of the reference genome to individual chromosomes, the mkrrgenome tool was used to scan the genome for MspI recognition sites (C′CGG) to retain only those fragments that fall in the specified size range of 20-200 bp. The resulting output was then converted into a region bed file format using a custom python script. For scripts and documentation, please visit the following GitHub repository https://github.com/VIBTOBIlab/BismarkPipeline (v1.0.0).

### Sequencing saturation curves

To calculate the sequencing saturation, we adopted the following strategy. For each *in silico* generated sample (e.g. an *in silico* mixture composed of 10% of tumour reads and 90% of healthy reads) we calculated the number of unique CpGs covered by at least 3 reads at the five different sequencing depths for RRBS samples (2M, 5M, 10M, 15M and 20M) and four sequencing depths for WGBS samples (62M, 183M, 310M and 620M). Subsequently, we used *scipy curve_fit function* to fit the following curve:

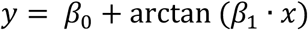

Where *y* corresponds to the number of CpGs covered by at least 3 reads, *x* corresponds to the number of reads/percentage of downsampling, the parameter *β*_0_ scales the maximum height (the asymptote) of the curve and *β*_1_ controls how fast the curve approaches the asymptote. We used the arctan function since it has an asymptotic growth similar to sequencing saturation. For x that tends to infinite values, we can compute the asymptote as the following:

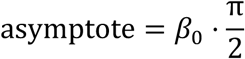

The sequencing saturation value (which ranges from 0 to 1) is then calculated as the number of unique CpGs with at least 3 reads observed, divided by the theoretical maximum number of CpGs, which corresponds to the asymptote of the fitted curve:

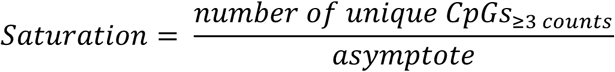

The scripts used to compute and plot the sequencing saturation curves are available at https://github.com/VIBTOBIlab/BenchmarkingAnalysis (v1.0.0).

### Benchmark execution

#### DMRs selection for reference-based deconvolution

To run the benchmarking of reference based tools, we used three different differentially methylated regions (DMRs) selection tools (Supplementary Table 1C): limma (with pre-defined regions), DMRfinder and wgbstools^28–30^. Both DMRfinder and wgbstools have built-in functionality to group individual CpGs into regions and test for statistically significant differential methylation. On contrary, for limma we used the hg19-RR region file in the DecoNFlow pipeline to merge the individual CpGs into regions using bedtools intersect function and finally tested these regions for differential methylation using limma workflow.

Regardless of the DMR detection tool used, we applied the following filtering criteria to the identified regions. For the WGBS-TT dataset, only regions containing at least three CpGs and with a minimum coverage of five counts per CpG were retained. For the RRBS-TT and RRBS-CL datasets, which provide higher CpG coverage, a stricter threshold of at least ten counts per CpG was applied. Next, for each tumour type, we identified the top 250 hypo-methylated regions in the tumour samples and the top 250 hypo-methylated regions in the healthy samples (500 regions in total), retaining only those with an adjusted p-value < 0.01 after Benjamini–Hochberg correction. When fewer than 250 DMRs per entity met this criterion, all DMRs with an adjusted p-value < 0.01 were retained. Finally, reference samples corresponding to each entity were averaged across regions to obtain a single representative value per entity. No filtering was applied to the *in silico* mixtures before deconvolution.

#### Regions selection for reference-free deconvolution

Before being deconvolved using the reference-free tools, the bulk samples were processed as follows. Individual CpGs with at least three counts were retained and grouped into regions using 2 different region files. One file is composed of regions after *in silico* MspI digestion of the hg19 genome, as described above. The second file is composed of regions covering CpGIslands in hg19 genome and was downloaded from UCSC Table Browser^48^ using the following query: *Clade: Mammal, Genome: Human, Assembly: hg19, Group: Regulation, Track: CpG Islands*. Both files can be found in the Supplementary Material.

#### DecoNFlow

To benchmark the 10 different reference-based and 2 reference-free deconvolution tools (Supplementary Table 1A^11,18–26^) across the three DMR selection approaches describe above and to assess their performance, we have implemented a reproducible and scalable Nextflow pipeline: DecoNFlow (Nextflow >= 23.04.0, https://github.com/VIBTOBIlab/DecoNFlow). This benchmarking was performed running DecoNFlow version v2.2.0. The pipeline can be run either locally or using Docker^49^ or Apptainer/Singularity^50^, allowing the users to run it on high-performance computing and/or Cloud infrastructures. Moreover, the pipeline allows to use customized configuration files for optimal execution on clusters or compute environments. Several publicly available config files can be found on nf-core (https://nf-co.re/configs/). Using the *--benchmark* flag, the pipeline will run all the methods simultaneously (given all the required input files). However, the pipeline can also run individual methods specifying the corresponding flag. DecoNFlow requires at least 8 CPUs and 16 GB of RAM to be run, making it possible to run it on a personal laptop as well. For a comprehensive description of the pipeline, look at the documentation provided in the GitHub repository (https://github.com/VIBTOBIlab/DecoNFlow). For more information about the parameters used to run the benchmark, see the parameter files present in Supplementary Material.

### Scalability of the tools

To compute the scalability of the tools, we used the following strategy. We selected 5, 25, 50 and 100 *in silico* mixtures from the NBL RRBS-CL dataset and we run each tool using sequentially 25, 50, 100, 250 and 500 DMRs. Second, we extracted the memory peak and the real time computed automatically by Nextflow for each individual deconvolution tool. Then, we averaged the time and memory for each combination to compute the average running time and memory usage. Finally, we computed the *Score*_*scalability*_ by min-max scaling the average running time and memory usage across the deconvolution tools. We ran this analysis allocating 8 CPUs and 16GB of memory to each deconvolution tool, the minimum required resources to run the pipeline.

### Evaluation metrics and missing values

Four metrics have been used in this study to evaluate the performance of the deconvolution tools: the inverse root-mean-squared error (1-*RMSE*), Spearman’s rank correlation coefficient (*SCC*), the area under the receiver operating characteristic curve (*AUC* − *ROC*) and the limit of detection (*LoD*). These four metrics are computed for each combination of DMR tool, sequencing depth and tumour type.

The *RMSE* is used to evaluate the error of the tools in predicting the tumoral fraction *tf_estimated_* and it is computed as follows:

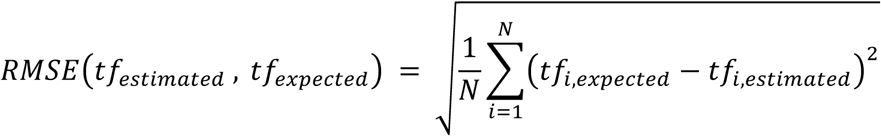

where *N* corresponds to the total number of samples present in each combination being analysed at observed tumoral fraction greater than 0. Since the *RMSE* is not a scale-free metric, it will increase at increasing observed tumoral fraction. This determines that the *RMSE* at high tumoral fraction (e.g. 0.25 and 0.5) will dominate the overall and advantage those tools that perform well uniquely at high tumoral fractions. To account for this bias when ranking the deconvolution tools, we compute the *RMSE* separately at each observed tumoral fraction within each combination, then min-max scale it across deconvolution tools and aggregate the individual min-max scaled *RMSE* at each tumoral fraction using the mean to get a unique *RMSE* per combination.

The SCC is used to evaluate the correlation between observed *X* and estimated *Y* tumoral fraction. To compute SCC, *X* and *Y* are converted to ranks *R*[*X*], *R*[*Y*]:

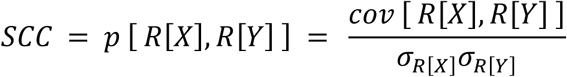

Where *p* is the conventional Pearson correlation coefficient operator applied to the rank variables, *cov* [ *R*[*X*], *R*[*Y*]] is the covariance of the rank variables and *σ*_*R*[*X*]_, *σ*_*R*[*Y*]_ are the standard deviations of the rank variables. The SCC is computed for each combination pooling all the observed tumour fractions together to get one value per combination.

The AUC value is used to measure the ability of a binary classifier to distinguish between positive and negative classes across all possible decision thresholds. In our context, the AUC value is used to measure the ability of deconvolution tools in predicting presence or absence of tumour at different tumour fractions. To compute the AUC, we compared samples with an observed tumour fraction of 0 (representing no tumor) to samples with tumour fractions greater than 0. For each observed tumour fraction, we binarized the labels as 0 (no tumour DNA present) and 1 (tumour DNA present). We then calculated the AUC for each pairwise comparison. As a last step, we aggregated the resulting AUC values by averaging them to obtain a single AUC per combination.

Finally, the LoD represents the lowest tumoral fraction that methods can detect. To determine the LoD for each tool, we identified the lowest tumour fraction (moving from higher to lower fractions) that remained statistically significant (adjusted p-value < 0.01) using an unpaired one-tailed Mann-Whitney U-test with BH multiple testing correction. In other words, the LoD corresponds to the last significant fraction before the first non-significant one. For each analysed combination the LoD of each deconvolution tool is computed.

To get a unique aggregated metric per combination, the metrics are min-max scaled across deconvolution tools at each combination analysed and aggregated into a final score, and then min-max scaled across tools:

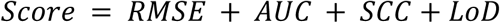

Subsequently, for each dataset we computed the average score across all combinations to get one unique score per dataset. Finally, to get a unique score per deconvolution tool across all the datasets, the individual scores per dataset were aggregated using the following weighted mean:

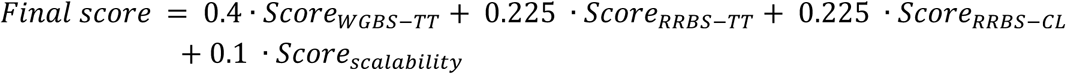

To avoid unbalancing the final score toward the RRBS sequencing method, we gave the weights 0.5, 0.25 and 0.25 to respectively WGBS-TT, RRBS-TT and RRBS-CL. Moreover, since all the tools run pretty fast (< 1min on average) and with a low memory requirement (< 1GB on average, excluding MeDeCom), we gave a weight of 0.1 to the scalability score to reduce its influence in the final ranking. Furthermore, although some of the tools produced NA values instead of predictions, we decided to just remove these values instead of penalizing the tools proportionally to the number of missing values, since only 7 and 2 observations (out of the 3960) were missing from EpiDISH and EpiSCORE, respectively (Supplementary Table 4)). To compute the final ranking, we ranked the tools from the highest to the lowest overall score following best practice guidelines^51^.

## Code availability

Pipeline used to preprocess the RRBS samples can be found at https://github.com/VIBTOBIlab/MethylSeq/tree/2.6.2. The nf-core/methylseq pipeline used to preprocess the WGBS samples can be found at https://nf-co.re/methylseq/2.6.0/ and https://github.com/nf-core/methylseq/tree/2.6.0. Pipeline used to perform in silico mixtures of WGBS dataset can be found at https://github.com/VIBTOBIlab/Sillymix/tree/1.1.0. DecoNFlow pipeline used to generate the benchmark results can be found at https://github.com/VIBTOBIlab/DecoNFlow. Scripts and documentation to generate an RR genome can be found here https://github.com/VIBTOBIlab/BismarkPipeline/tree/v1.0.0. Scripts necessary to reproduce the results and generate the plots can be found at https://github.com/VIBTOBIlab/BenchmarkingAnalysis (v1.0.0). The R shiny app and the source code containing interactive plots can be found here at https://github.com/VIBTOBIlab/Shiny-DNAmBenchmarking (v1.0.0).

## Data availability

WGBS tumoral samples were downloaded from TCGA using the corresponding manifests present at the GDC Data Release 36.0 (https://docs.gdc.cancer.gov/Data/Release_Notes/Data_Release_Notes/#data-release-360) controlled and maintained by dbGaP. The 12 WGBS healthy cfDNA samples can be found in NCBI GEO under accession number <u>GSE164600.</u> Healthy reference RRBS plasma samples used for the RRBS-TT and RRBS-CL datasets were downloaded from the EGA database (EGAD50000000734). CRC, LC and PC tumour samples can be found in NCBI GEO under accession number <u>GSE79211</u>. Data from the NBL cell lines used for both the in silico mixtures and the reference samples of the RRBS-CL dataset is being submitted to EGA. Likewise, the healthy pooled cfDNA used for generating the in silico mixtures in both the RRBS-TT and RRBS-CL datasets is also being submitted to EGA.

## Acknowledgments

We acknowledge the use and support of the VIB Data Core facility. We also thank the HPC-UGent team for their support. This project was supported by Kom op tegen Kanker [11559], the European Research Council (ERC) under the European Union’s (Horizon 2020 research and innovation programme) [101044243] and the Fonds Wetenschappelijk Onderzoek (FWO) PhD Fellowship [1126825N].

