## Supplementary Figures for "A benchmark of DNA methylation deconvolution methods for tumoral fraction estimation using DecoNFlow"

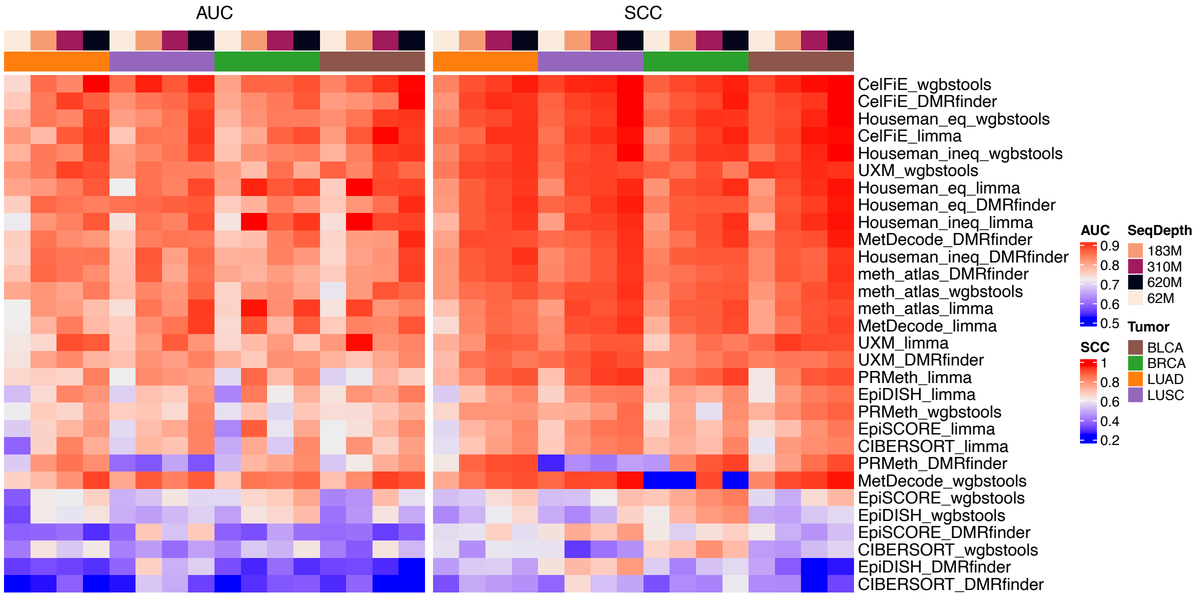


**Supplementary Figure 1.** Heatmap showing the raw AUC and SCC on the WGBS-TT dataset for each deconvolution-DMR method combination. The SCC was computed across all different expected tumoral fractions, while the AUC was computed as the average of the AUC-ROC values at each individual tumoral fraction. We computed the row mean and ordered the rows by descending order.

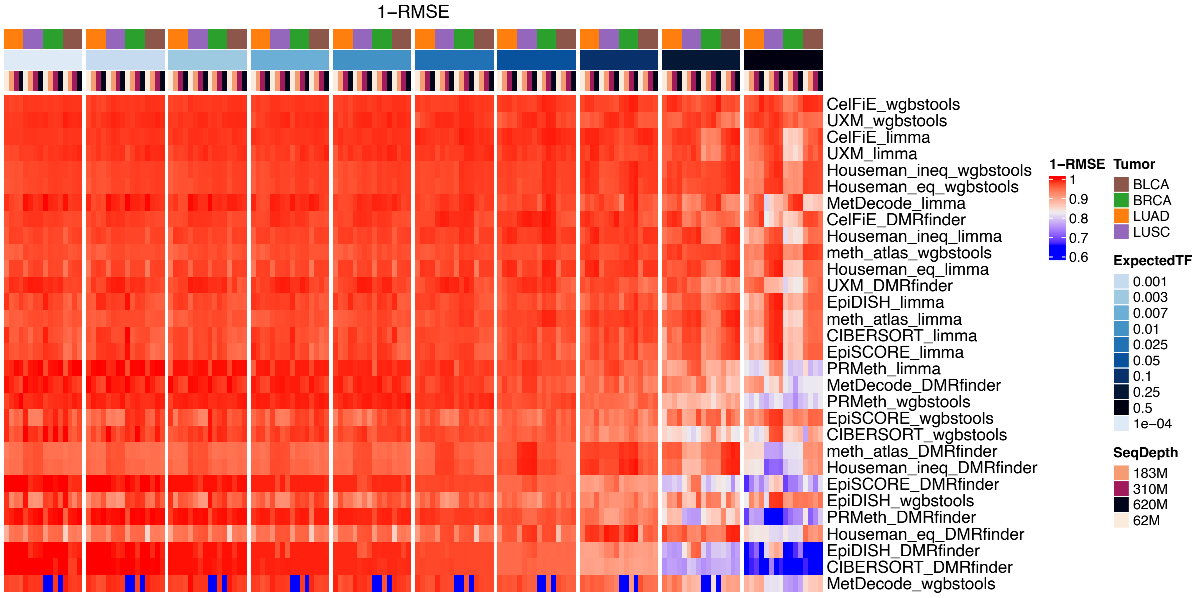


**Supplementary Figure 2.** Heatmap showing the raw RMSE on the WGBS-TT dataset computed over the 10 sample replicates at each expected tumoral fraction, tumor type, sequencing depth and combination of deconvolution tool and DMR tool. The rows were ordered by descending 1-RMSE value.

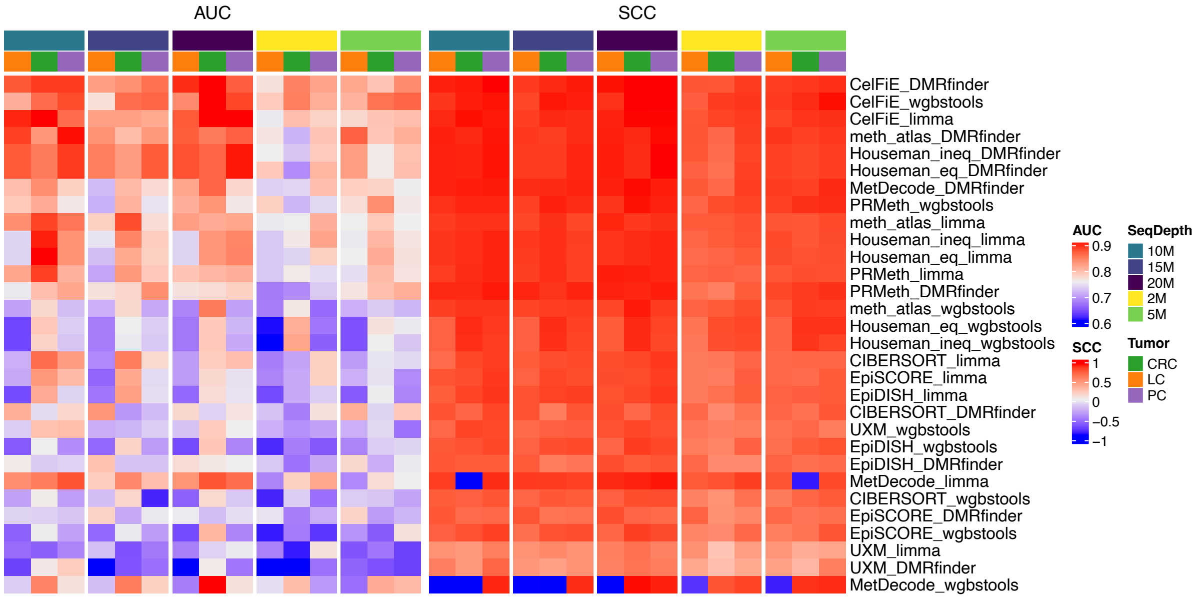


**Supplementary Figure 3.** Heatmap showing the raw AUC and SCC on the RRBS-TT dataset. The SCC was computed across all different expected tumoral fractions, while the AUC was computed as the averaging the AUC-ROC values at each individual tumoral fraction. The rows were ordered by descending row mean.

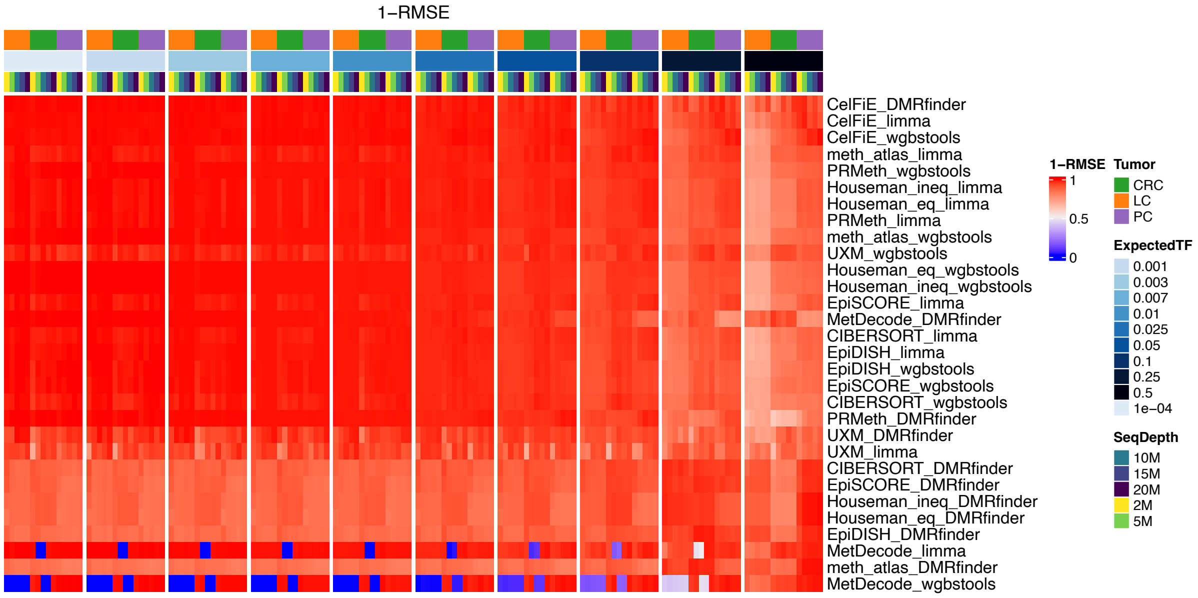


**Supplementary Figure 4.** Heatmap showing the raw RMSE on the RRBS-TT dataset computed over the 10 sample replicates at each expected tumoral fraction, tumor type, sequencing depth and combination of deconvolution tool and DMR tool. The rows were ordered by descending 1-RMSE value.

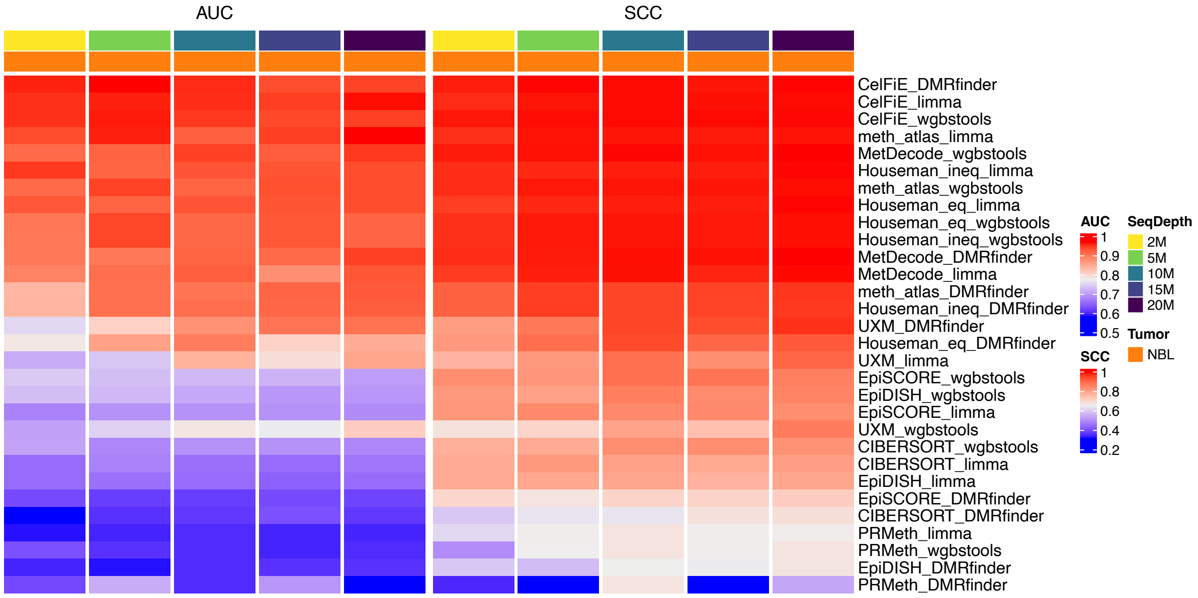


**Supplementary Figure 5.** Heatmap showing the raw AUC and SCC on the RRBS-CL dataset. The SCC was computed across all different expected tumoral fractions, while the AUC was computed as the averaging the AUC-ROC values at each individual tumoral fraction. To order the rows, we computed the row mean and ordered by descending order.

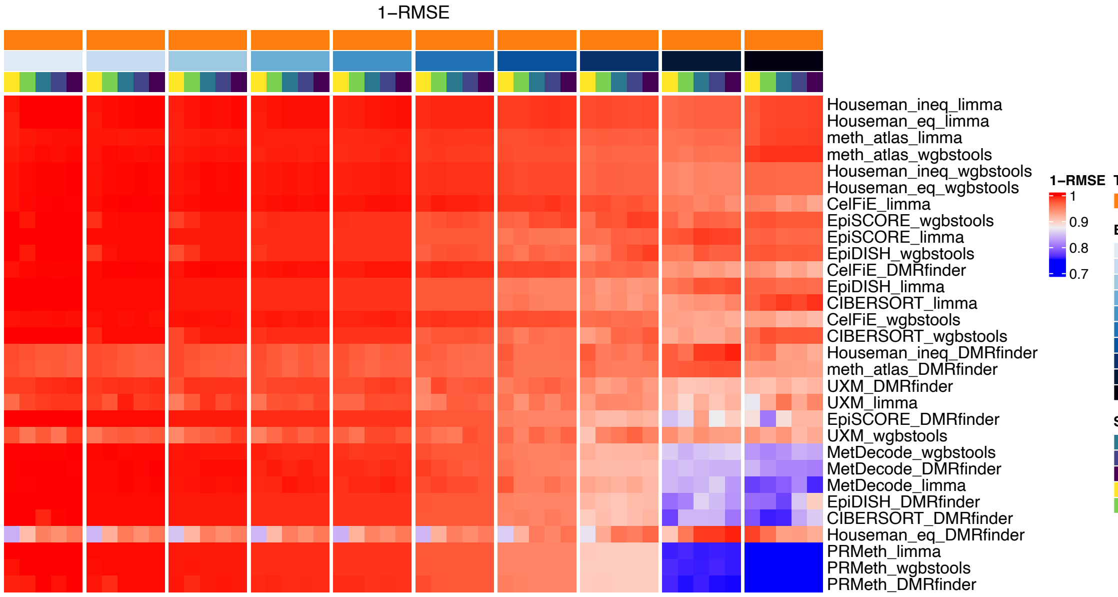


**Supplementary Figure 6.** Heatmap showing the raw RMSE on the RRBS-CL dataset computed over the 10 sample replicates at each expected tumoral fraction, tumor type, sequencing depth and combination of deconvolution tool and DMR tool. The rows were ordered them by descending 1-RMSE.

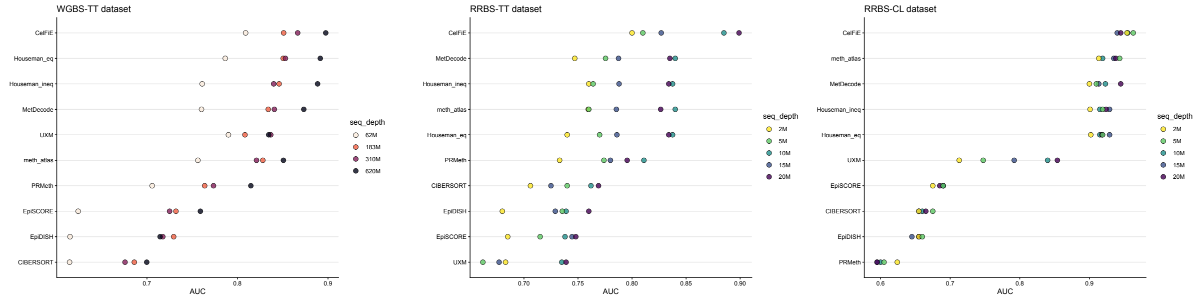


**Supplementary Figure 7.** Dotplots for WGBS-TT, RRBS-TT and RRBS-CL datasets showing the median AUC-ROC values for each sequencing depth. The deconvolution tools are then ordered from the higher to the lower mean AUC-ROC values across sequencing depths.

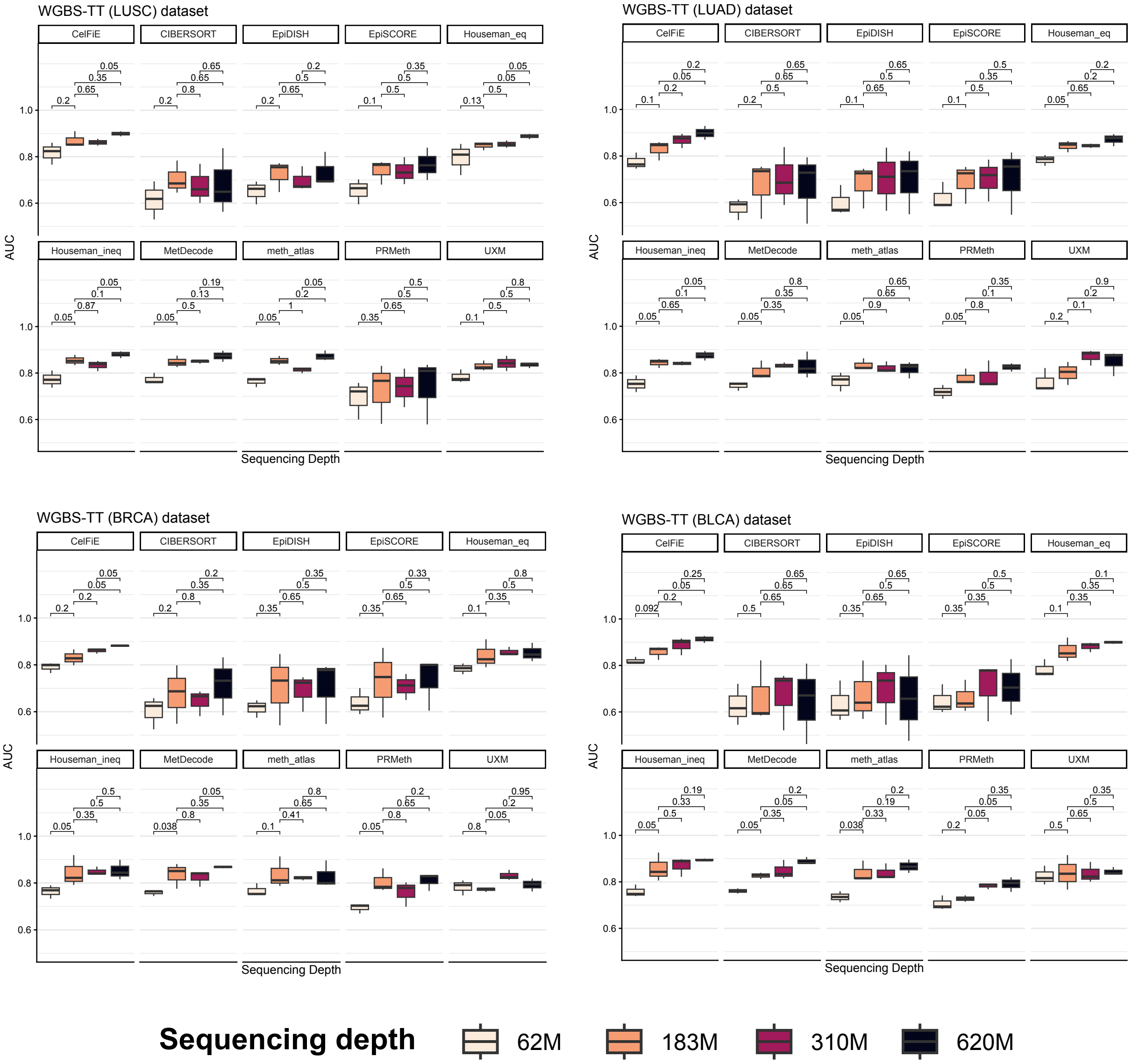


**Supplementary Figure 8.** Boxplots of AUC-ROC values in WGBS-TT dataset at each sequencing depth divided by deconvolution tool. An unpaired Mann-Whitney U test (with BH correction) was performed at increasing sequencing depths to test whether the increase in AUC-ROC at increased sequencing depth was statistically significant or not.

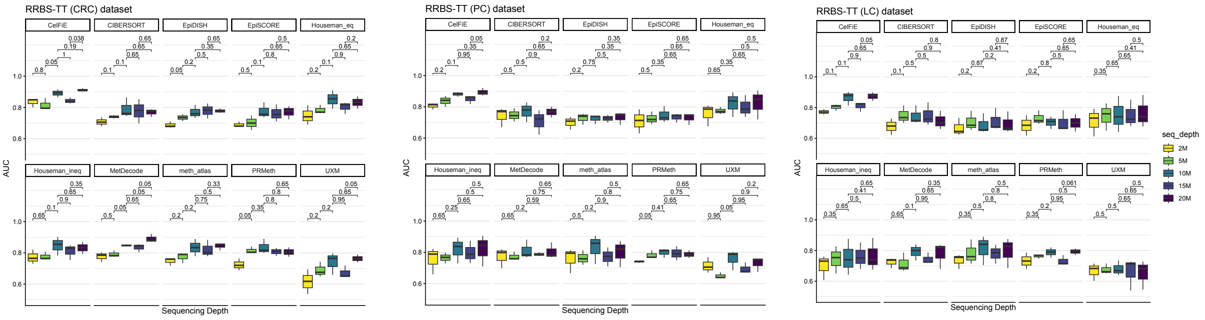


**Supplementary Figure 9.** Boxplots of AUC-ROC values in RRBS-TT dataset at each sequencing depth divided by deconvolution tool. An unpaired Mann-Whitney U test (with BH correction) was performed at increasing sequencing depths to test whether the increase in AUC-ROC at increased sequencing depth was statistically significant or not.

**A**

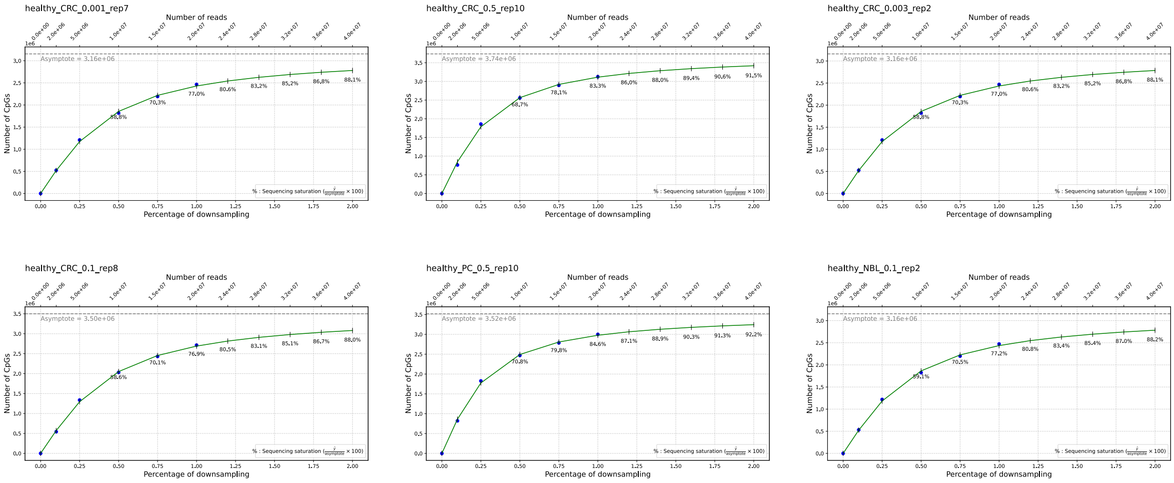


**B**

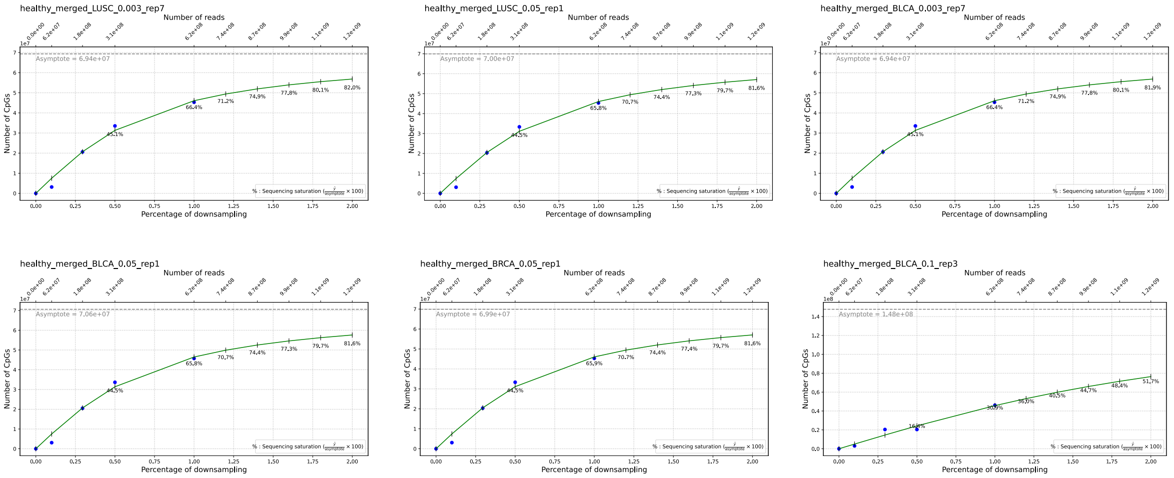


**Supplementary Figure 10. (A)** Sequencing saturation curves of six RRBS-based sequencing samples: NBL 10% tumor fraction, CRC 10% tumor fraction, CRC 50% tumor fraction and PC 50% tumor fraction, CRC at 0.3% and 0.1%. The plots show the saturation (elbow of the curve) reached at 10M of reads, correponding to the value at which increasing the number of reads does not result in a linear increase of unique CpG (covered by at least 3 reads) discovered. **(B)** Sequencing saturation curves of six WGBS-base sequencing samples: BRCA 5%, BLCA 5%, LUSC 5%, BLCA 1%, LUSC 0.3% and BLCA 0.3%. In WGBS sequencing saturation plots sequencing saturation is lower and plateau still not reached, suggesting that increasing the number of reads would linearly increase unique CpGs discovered.

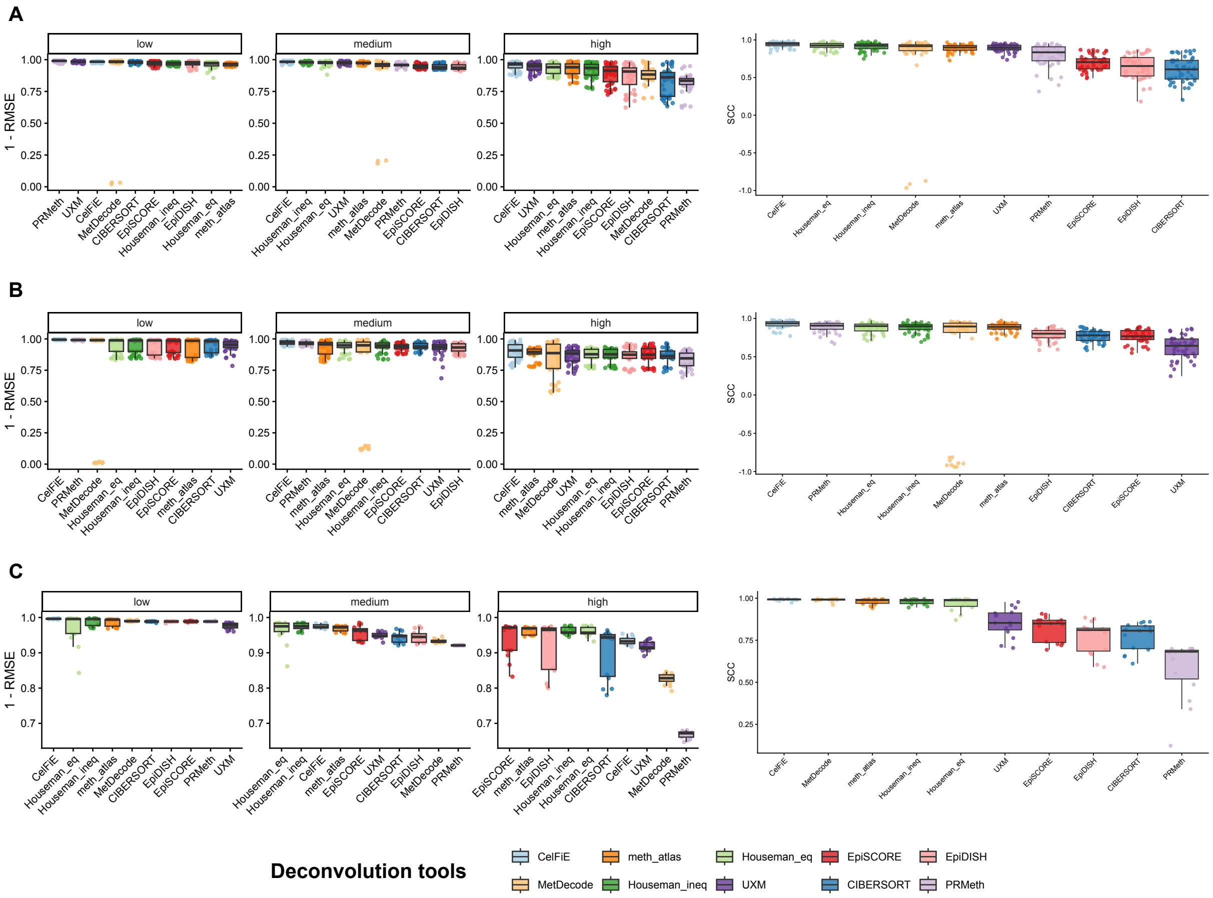


**Supplementary Figure 11.** Preciseness of the deconvolution tools evaluated according to the SCC and 1-RMSE. The 1-RMSE was plotted at three different expected tumoral fractions to avoid skewing thr distribution to the higher values: low (≤2.5%), medium (2.5% to 25% excluded), and high (≥25%). Each dataset is plotted separately for a more detailed information: A) WGBS-TT; B) RRBS-TT and C) RRBS-CL.

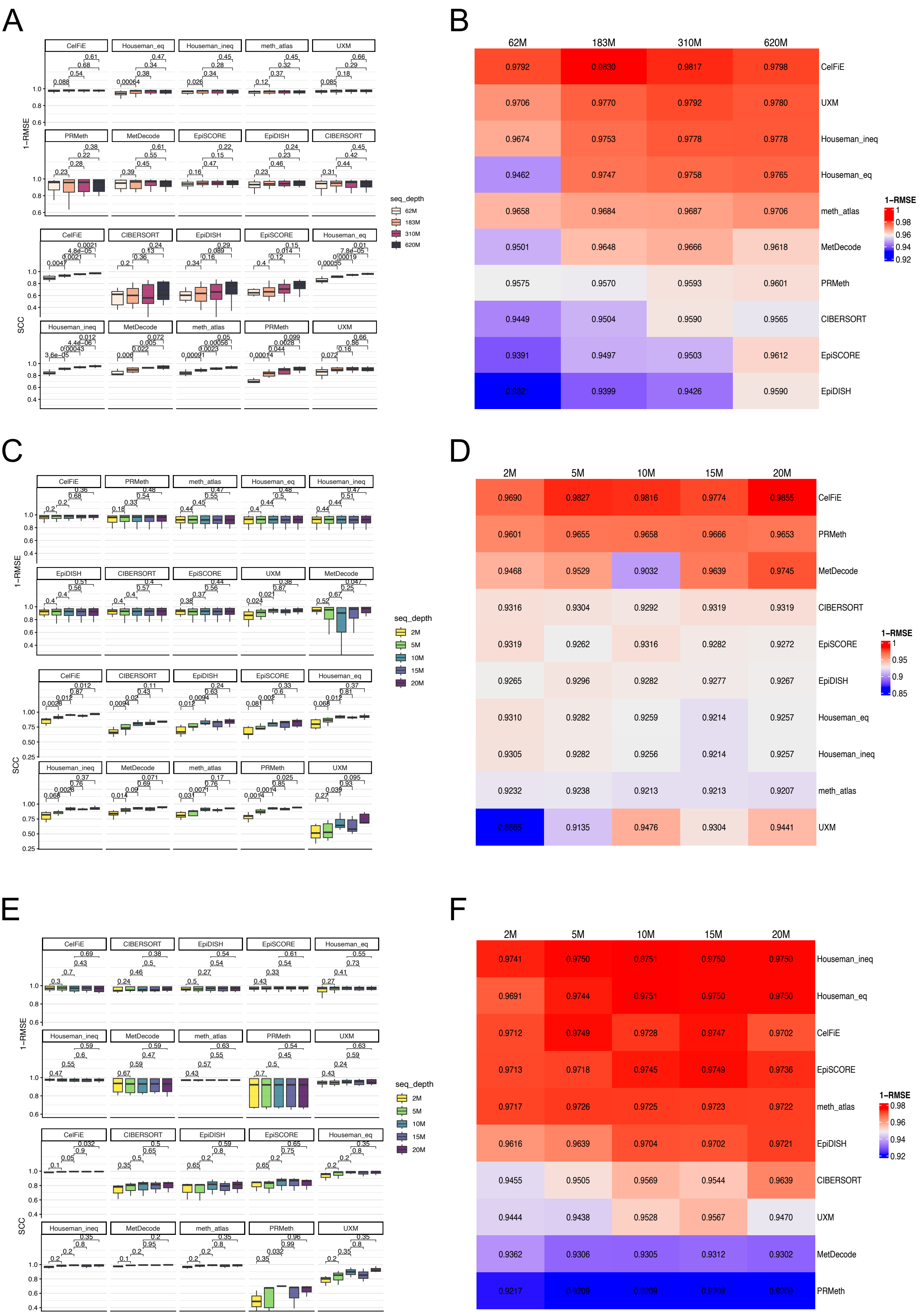


**Supplementary Figure 12.** Boxplots of 1-RMSE and SCC values in WGBS-TT **(A),** RRBS-TT **(C)** and RRBS-CL **(E)** dataset at each sequencing depth divided by deconvolution tool. An unpaired Mann-Whitney U test (with BH correction) was performed at increasing sequencing depths to test whether the increase in 1-RMSE and SCC at increased sequencing depth was statistically significant or not. Subplots **B, D** and **F** show raw median 1-RMSE values at each sequencing depth.

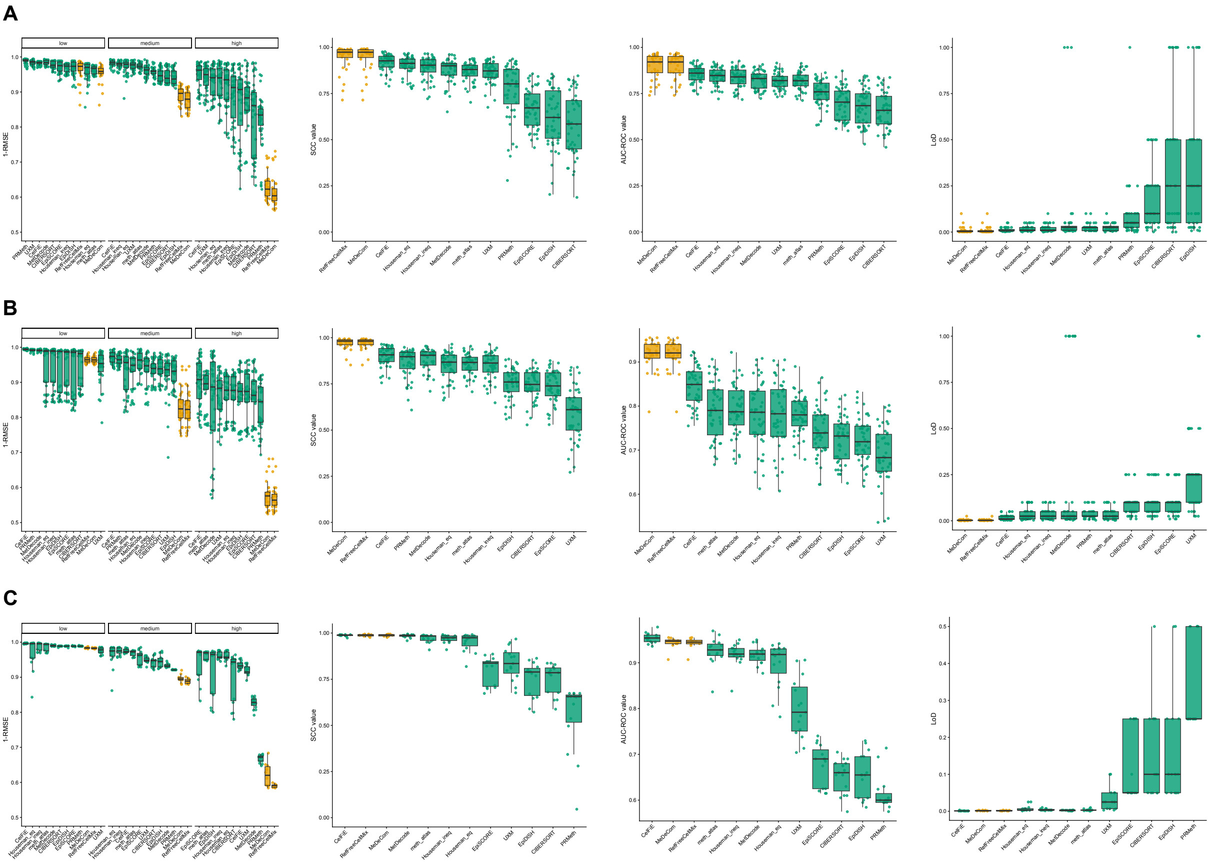


**Supplementary Figure 13.** From left to right**,** 1-RMSE, SCC, AUC-ROC and LoD score distributions shown separately for each WGBS-TT (A), RRBS-TT (B) and RRBS-CL (C) datasets.

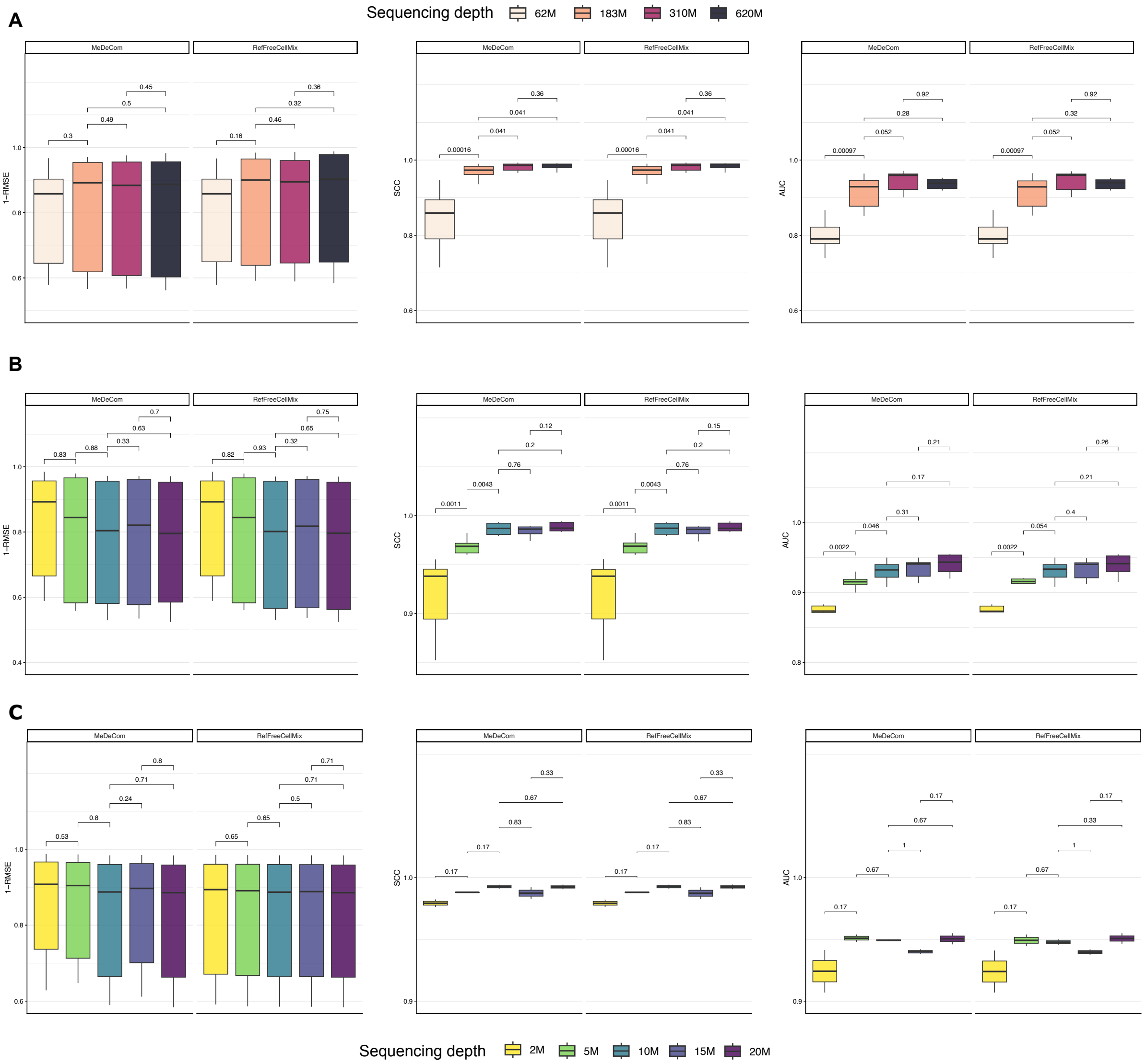


**Supplementary Figure 14.** MeDeCom and RefFreeCellMix 1-RMSE, SCC and AUC distributions at each sequencing depth for tools for the WGBS-TT **(A),** RRBS-TT **(B)** and RRBS-CL **(C)** datasets. An unpaired Mann-Whitney U test (with BH correction) was performed for eeach metric at increasing sequencing depths to test whether the increase at increased sequencing depth was statistically significant.
