## Supplementary Methods for "A benchmark of DNA methylation deconvolution methods for tumoral fraction estimation using DecoNFlow"

### Human neuroblastoma cell lines

In total, we collected DNA from 10 neuroblastoma cell lines with different phenotypes and genotypes that were used for reference building or for the *in silico* dataset generation (Table 1). The CLB-GA cell line (for *in silico* mixture generation) was grown in RPMI-1640 medium supplemented with 10% FCS, L-glutamine (2 mM), and HEPES (25 mM) and Penicillin (100 IU/ml), and Streptomycin (100 IU/ml). Cells were cultured at 37°C in a 5% CO2 humidified environment. Mycoplasma testing (Lonza) and short tandem repeat genotyping were regularly performed.

**Table 1.** Overview cell lines used in the benchmarking study. A: Adrenergic phenotype; M: Mesenchymal phenotype; Mi: Mixed phenotype.

| Cell line | A/M/Mi | ALK mutation | MYCN amplified | Origin | Used for NBL reference | Used for in silico mixtures |
|---|---|---|---|---|---|---|
| CLB-GA | A | R1275Q | No | Centre Leon Bérard, Lyon (FRANCE) | No | Yes |
| GI-M-EN | M | wt | No | Amsterdam Medical Center (NETHERLANDS) | Yes | No |
| Kelly | A | F1174L | Yes | Sigma-Aldrich (BELGIUM) | Yes | No |
| NB1 | A | wt | Yes | Amsterdam Medical Center (NETHERLANDS) | Yes | No |
| SH-SY5Y | A | F1174L | No | VIB-UGent , Center for Medical Biotechnology (BELGIUM) | Yes | No |
| SHEP | M | F1174L | No | University of San Francisco (UNITED STATES OF AMERICA) | Yes | No |
| SK-N-AS | Mi | wt | No | American Type Culture Collection (ATCC) (UNITED KINGDOM) | Yes | No |
| SK-N-BE(2)-C | A | wt | Yes | Northern Institute for Cancer Research (UNITED KINGDOM) | Yes | No |
| SK-N-FI | A | wt | No | Amsterdam Medical Center (NETHERLANDS) | Yes | No |
| SK-N-SH | Mi | F1174L | No | Amsterdam Medical Center (NETHERLANDS) | Yes | No |

### MNase digestion

Micrococcal nuclease (MNase) enzymatic gDNA fragmentation was performed within the CLB-GA cell line using aliquots of 7,5 x $10^6$ cells. Cells were pelleted for 5 minutes at 1200 rcf at 4°C and rinsed twice in ice cold PBS. Next, cells were permeabilized with ice-cold 1% Triton-PBS for 5 minutes, followed by 1 minute of gentle vortexing and washed twice in MNase digestion buffer (1 mM Tris-base, 5 mM CaCl2, 5 mM 2-mercaptoethanol, 0.1g/L BSA, supplemented with EDTA-free protease inhibitors (cOmplete™ Mini, EDTA-free Protease Inhibitor Cocktail, Roche)), after which they were resuspended in 100µl MNase digestion buffer. The nuclei mixture was preheated to 37°C, after which it was supplemented an MNase gradient (New England Biolabs, M0247S) over four tubes (1200U – 1400U – 1600U – 1800U) and incubated at 37°C for 15 minutes. Digestion was halted by adding stop buffer (5X solution, 250 mM EDTA, 250 mM EGTA). The cells were diluted with PBS to an input volume of 2 ml, and fragmented DNA (artificial cfDNA, arti-cfDNA) was isolated using the Maxwell RSC LV ccfDNA kit (Promega, AS1840) according to manufactures protocol. The length profile was evaluated via TapeStation cell-free DNA ScreenTape platform (Agilent, 5067-5630 & 5067-5631) and DNA concentration was measured using the Quant-iT dsDNA High-Sensitivity (HS) Assay Kit (Thermo Scientific, Q33120). Samples were selected when they contained at least 80% of cfDNA called by the TapeStation software (fragments between 70 and 700 bp), and when they showed the profile of mono-nucleosomes and a smaller peak of di-nucleosomes.

### cfDNA isolation from healthy donors

Sample collection was approved by the ethics committee of Ghent University Hospital (Belgian Registration number B670201733701) and written informed consent was obtained from 15 healthy donors. Venous blood from the healthy volunteers was collected in at least 2 BD Vacutainer Plastic K2EDTA tubes per volunteer (EDTA; Becton Dickinson and Company, 367525) from the antecubital vein after disinfection with 2% chlorhexidine in 70% alcohol (BD Vacutainer Push Button Blood Collection Set (BD, 367326). Plasma isolation was immediately performed by centrifugation of the blood samples at 1900g for 10 min to obtain platelet-rich plasma (PRP), followed by a second identical centrifugation to remove platelets and white blood cells from the PRP. Plasma was stored at -80°C until further processing. 7.5mL of plasma was thawed on ice, and cfDNA was extracted using the Maxwell RSC LV ccfDNA kit (Promega, AS1840) according to the manufacturer's instructions. Quality control of the cfDNA traces included to check the absence of high molecular weight DNA and was performed using the TapeStation cell-free DNA ScreenTape platform (Agilent, 5067-5630 & 5067-5631). DNA concentration was measured using the Quant-iT dsDNA High-Sensitivity (HS) Assay Kit (Thermo Scientific, Q33120). To obtain sufficient cfDNA from healthy volunteers for downstream processing, cfDNA passing QC was pooled across donors.

### RRBS library construction

RRBS library construction was performed as described in De Koker et al[1]. and Van Paemel *et al*[2]. In short, up to 10ng DNA was used for rSAP dephosphorylation, MspI digest and A-tailing of the DNA. Subsequently, hairpin adapters are ligated to the digested DNA, and fragments that were not completely digested with MspI (dephosphorylated ends) are degraded with exonuclease before adapter opening using USER enzyme. Finally, MspI enriched DNA is bisulfite converted before library amplification. The libraries were visualized with the NGS Fragment Kit (1-6000bp) (Agilent, DNF-473-0500) and quantified using the KAPA Library Quantification Kits (Roche, KK4854). The libraries were equimolarly pooled and were sequenced on a NovaSeq 6000 instrument with a NovaSeq SP kit (paired-end, 2×50 cycles), using 3% phiX and a loading concentration between 500 and 750 nM.
